# Mapping of CD8 T-cell recognition to latent EBV infection and neuroantigens reveals HLA-specific depletion of T-cell responses in multiple sclerosis

**DOI:** 10.1101/2025.10.23.682742

**Authors:** Nikolaj P Kristensen, Natasja W Ettienne, Lasse F Voss, Amalie K Bentzen, Marina R von Essen, Tripti Tamhane, Finn Sellebjerg, Sine R Hadrup

## Abstract

Multiple sclerosis (MS) is a complex autoimmune disease with an unclear contribution from antigen-specific CD8 T cells. To enumerate adaptive immune responses associated with MS and to identify MS-related epitopes recognized by CD8 T cells, we evaluated T-cell recognition of 1,158 predicted major histocompatibility complex (MHC)-binding peptides derived from latent EBV antigens (laEBVs) and selected neuroantigens (NAgs) using DNA barcode-labelled pMHC multimers. HLA-B*07:02+ individuals living with relapsing-remitting MS (RRMS) and progressive MS (PMS) were characterized by a relative reduction in EBNA6-specific and MAG-specific CD8 T cells compared to healthy donors. Furthermore, both RRMS and PMS patients exhibited a delayed expansion of CMV- and laEBV-specific CD8 T cells compared to healthy donors indicating an age-specific interaction with disease. Female RRMS patients were moreover characterized by a specific reduction of laEBV-specific CD8 T cells compared to male RRMS patients. NAg-specific CD8 T cells recognized heterogeneous peptides, were of low cellular frequency in peripheral blood, and exhibited a circulating, naive-like, anergic phenotype. In contrast, virus-specific CD8 T cells were characterized by high-frequency cellular responses with an effector-like, tissue-homing phenotype. These immunophenotypes were largely shared between MS and healthy donors.

Collectively, our data provide novel insights into CD8 T-cell recognition of both NAg and laEBV antigens in MS and constitute a key reference map of epitopes for further immunomonitoring of EBV- and self-specific CD8 T cells in health and disease.

## INTRODUCTION

Multiple Sclerosis (MS) is a demyelinating disease of the central nervous system that often manifests in early adulthood and has a higher incidence in women (McGinley et al., 2021). The pathology is characterized by immune cell infiltration, progressive demyelination, loss of oligodendrocytes and axonal damage in the brain and spinal cord (Kuhlmann et al., 2023; Lassmann, 2018). The disease can be stratified into two subgroups based on clinical symptoms: Relapsing-remitting MS (RRMS) and progressive MS (PMS). PMS can be further subdivided into primary PMS or secondary PMS, with or without a prior RRMS diagnosis, respectively (Lublin et al., 2014).

Many environmental and genetic factors are associated with MS (Attfield et al., 2022; International Multiple Sclerosis Genetics Consortium., 2019). Among these, Epstein-Barr Virus (EBV) infection (Bjornevik et al., 2022; Thacker et al., 2006) and the major histocompatibility complex (MHC) class II-encoding HLA-DRB1*15:01 represent the strongest independent risk factors for the development of MS (International Multiple Sclerosis Genetics Consortium., 2019; Moutsianas et al., 2015). Multiple HLA class I alleles (HLA-A*02:01, B*38:01, B44:02 and B*55:01) are furthermore associated with protection from MS (Moutsianas et al., 2015).

T cells are generally thought to contribute to disease pathology due to their interaction with peptides presented by MHC class I and II. This is substantiated by (i) the capacity of neuroantigen(NAg)-specific T cells to drive MS-like symptoms in the murine experimental autoimmune encephalomyelitis (EAE) model (Glatigny and Bettelli, 2018), (ii) the increased presence of brain-infiltrating T cells in MS patients compared to non-inflammatory controls (Machado-Santos et al., 2018; van Nierop et al., 2017), and (iii) the accumulation of both perivascular and parenchymal brain-infiltrating T cells in active MS lesions compared to normal-appearing white matter (Fransen et al., 2020; Veroni et al., 2018).

Cytotoxic CD8 T cells are especially of interest in humans, because they outnumber other lymphocyte subsets in the early immune cell infiltrate of white-matter lesions (Fransen et al., 2020). As such, CD8 T cells may either directly facilitate tissue damage, or fail to prevent tissue damage from EBV-infected B cells in some capacity (Hassani et al., 2018; Moreno et al., 2018; Serafini et al., 2007; Veroni et al., 2018). This is further supported by the finding, that latent EBV infection of B cells in MS patients is poorly controlled (Soldan et al., 2024), possibly due to efficient immune evasion of cytotoxic lymphocytes by infected B cells through viral interleukin-10 (Bejarano and Masucci, 1998; Suzuki et al., 1995), inhibitory HLA-E ligands (Vietzen et al., 2023) and expression of inhibitory PD-L1 on EBV infected B cells (Serafini et al., 2024). The presence of strong immune evasion mechanisms coincides with an initial high frequency of EBV-reactive CD8 T cells during each autoimmune attack (Angelini et al., 2013; Jilek et al., 2008; Pender et al., 2017) followed by a decrease in the observed frequency of EBV-reactive T cells to below the level of healthy controls while in remission or in progressive stages of MS (Pender et al., 2017). Overall, this indicates that CD8 T cells likely fail to control latent EBV infection leading to persistent infection of B cells and retention of autoimmune B-cell responses, which may further exacerbate tissue damage through molecular mimicry (Lanz et al., 2022; Vietzen et al., 2023).

Previous reports monitoring EBV-reactive T cells among MS patients typically employed stimulation-based culture assays using either viral lysates, peptide pools or lymphoblastoid cell lines (LCLs) to enumerate cytokine-releasing CD8 T cells responding to EBV or EBV-derived antigens. Although useful to define antigen-reactivity, they do not readily allow for strict epitope-wise resolution and enumeration of EBV-specific CD8 T cells – nor do they allow for efficient discovery of EBV-derived epitopes. It is furthermore hypothesized that MS patients harbor antigen-specific CD8 T cells that recognize NAg-derived epitopes, possibly through molecular mimicry (Wucherpfennig and Strominger, 1995), and that such recognition should be absent in healthy individuals. We therefore tested more than one thousand predicted NAg and laEBV candidate epitopes using pMHC multimers on peripheral blood samples of both MS patients and healthy donors. Furthermore, we compared the cellular frequency of antigen-specific CD8 T cells, epitope recognition prevalence, and immunophenotypic profiles between patients and healthy donors to better understand how each population of antigen-specific CD8 T cells (NAg vs virus-specific) were related to MS.

We focused our investigation on immune responses restricted to 4 common human leukocyte antigen (HLA) alleles HLA-A*01:01, A*02:01, B*07:02 and B*08:01 across 4 laEBV antigens namely EBV nuclear antigen (EBNA) 3, EBNA4, EBNA6 and latent membrane protein 2 (LMP2). Furthermore, we included peptides derived from 5 NAgs previously shown to be recognized by T cells patients with MS including myelin-associated glycoprotein (MAG) (Sabatino et al., 2019; Tsuchida et al., 1994), myelin basic protein (MBP) (Jurewicz et al., 1998; Sabatino et al., 2019), myelin oligodendrocyte glycoprotein (MOG)(Sabatino et al., 2019; Tsuchida et al., 1994), myelin proteolipid protein (PLP)(Honma et al., 1997; Sabatino et al., 2019; Tsuchida et al., 1994), and transaldolase (TALDO1)(Niland et al., 2005).

We also included a panel of citrullinated NAg-derived peptides (CitNAgs), in which one or more central arginines were deaminated – a modification hypothesized to contribute to the loss of T-cell tolerance in MS (Teresa et al., 2023).

We observed that patients living with MS largely recognize the same set of epitopes as heathy donors. Nevertheless, we demonstrated a significant global reduction in the cellular frequency of naive-like/anergic NAg-specific CD8 T cells in peripheral blood of MS patients compared to healthy donors, which was further pronounced in context of HLA-B*07:02, specifically. We furthermore show that EBNA6-specific CD8 T cells in context of HLA-B*07:02 are specifically reduced in RRMS patients supporting previous claims of HLA-B*07:02-specific defects in EBV control (Agostini et al., 2018; Jilek et al., 2012). Finally, we observed weak age- and sex-dependent interactions, whereby MS patients exhibit altered or delayed development of laEBV- and NAg/CitNAg-specific CD8 T cell responses over time, with females showing higher NAg-specific but lower laEBV-specific CD8 T cell frequencies – particularly in the context of HLA-A*02:01 and B*07:02—compared to males.

Based on our broad epitope reference map and associated immune monitoring data, we speculate that EBV-specific CD8 T cells recognizing latent epitopes likely contribute to protection in context of specific HLA alleles, where a lack (or delayed development) thereof is associated with MS.

## RESULTS

### Neuroantigen-specific CD8 T cells are exceedingly rare in peripheral blood

We investigated CD8 T-cell recognition of 5 candidate NAgs and 4 known latent EBV (laEBV) antigens in peripheral blood of 20 healthy donors (HD), 19 patients living with RRMS and 19 patients living with PMS. 7 patients among PMS donors were diagnosed with primary PMS and 12 donors were diagnosed secondary PMS. PMS donors were generally older (52.1±8.7 years) and with higher Expanded Disability Status Scale (EDSS) scores (5.84±0.80 EDSS) than RRMS patients (41.6±10.4 years, 2.97±1.15 EDSS) (Table 1). Both RRMS and PMS donors were generally older than healthy donors (HD) (33.2±10.4 years).

**Table 1.**

|  | HD | RRMS | PMS |
| --- | --- | --- | --- |
| Donors (n) | 20 | 19 | 19<sup>†</sup> |
| Sex (F / M) | 15 / 5 | 12 / 7 | 9 / 10 |
| Age (±SD, years) | 33.2 (10.3) | 41.6 (10.4) | 52.1 (8.7) |
| Duration (±SD, years) | NA | 11.8 (6.0) | 10.1 (7.5) |
| EDSS (±SD) | NA | 2.97 (1.15) | 5.84 (0.80) |

*Age, duration and EDSS values are displayed as mean ± SD. EDSS, Expanded Disability Status Scale. Duration, disease duration. NA, not annotated. <sup>†</sup>7 primary progressive MS (PPMS) and 12 secondary progressive MS (SPMS).*

Differences in disease duration and sex ratios between cohort groups were negligible (Table 1). The distribution of the selected HLA Class I alleles (A*01:01, A*02:01, B*07:02, and B*08:01) were furthermore similar across the three cohort groups, although the PMS group featured slightly more B*07:02+ individuals than healthy donors (Supplemental table 1). HLA-DRB1*15:01 was significantly more represented among both patient groups compared to the healthy donor group (Supplemental table 1), further supporting this as a risk allele.

We predicted candidate peptides binding to HLA-A*01:01, A*02:01, B*07:02, or B*08:01 using netMHCcons1.1 (Karosiene et al., 2012)(Figure 1A) with native MAG, MBP, MOG, PLP and TALDO1, as well as Epstein-Barr nuclear antigen 3, 4 and 6 (EBNA3, 4 and 6) along with latent membrane protein 2 (LMP2) as input. Candidate NAgs were furthermore used as input for prediction of citrullinated, major histocompatibility complex (MHC)-binding peptides. We furthermore included a small set of 14 immunodominant peptides from influenza A (FLU-A), acute EBV infection, human cytomegalovirus (CMV) and human immunodeficiency virus (HIV) infection as positive and negative controls, which we collectively termed “CEF” (Supplemental table 2). The final library yielded 413, 475, 268 candidate peptides from laEBV antigens, native- and citrullinated neuroantigens (CitNAg), respectively (Figure 1A). B*07:02-binding peptides were the most numerous among predicted laEBV and CitNAg candidates with 195 and 124 peptides respectively, while A*02:01-binding peptides were the most numerous among native NAg-derived candidates with 224 peptides (Figure 1B).

**Figure 1:**
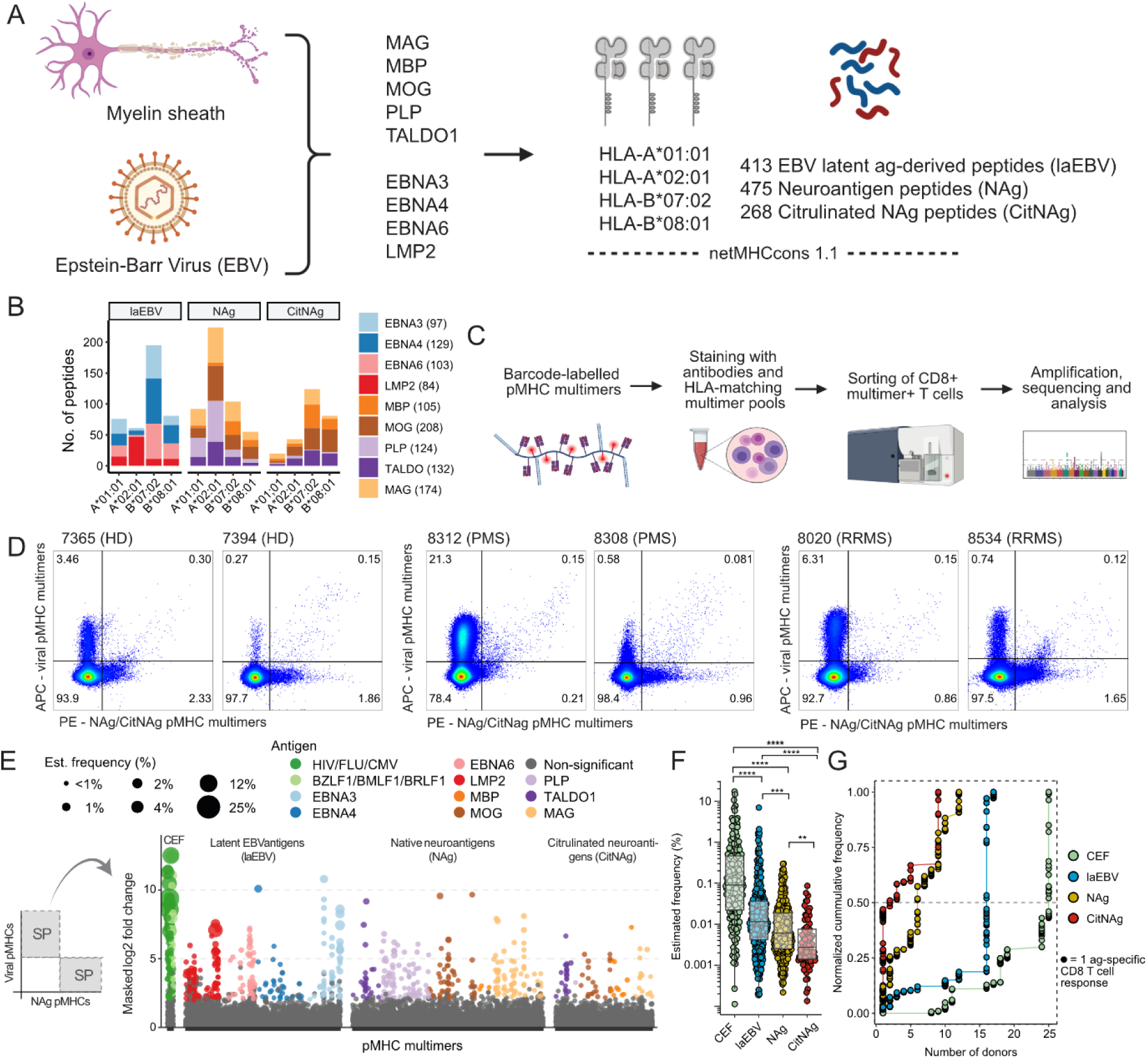
Identification of NAg and laEBV antigen-specific CD8 T cells in MS. A) BioRender illustration of antigen selection and peptide prediction. B) Multimer-library size across HLA and antigen groups. C) BioRender illustrated workflow using barcode-labelled multimers for discovery of antigen-specific CD8 T cells. D) Representative sorting plots. E) Epitope recognition landscape showing all 58 donors and enrichment scores (log2 fold change) from amplification of multimer-attached DNA-barcodes. Negative enrichment scores were masked i.e. set to 0. Grey represents non-enriched / non-significant DNA-barcodes. The order of sequences corresponds to the starting position for the given peptide within the antigen. F) Estimated frequency of epitope-specific CD8 T cells. Each dot indicates individual epitope-specific immune response. Colors indicate the number of independent donors recognizing the same epitope. G) Normalized cumulative cellular frequency as a function of the number of donors that recognize a given epitope. P-values were calculated using Barracoda (see materials and methods) for E, and Dunn’s test for F followed by adjustment for multiple hypothesis testing using the Benjamini-Hochberg approach. **, ***, and **** represent adjusted p-values lower than or equal to 0.01, 0.001, and 0.0001 respectively. Subfigures A and C were created in BioRender. Hadrup, S. (2025) https://BioRender.com/2tsw9dy.

Libraries of DNA barcode-labelled peptide-MHC (pMHC) dextran multimers were assembled for each peptide as previously described (Bentzen et al., 2016) and used as multimer pools for sorting pMHC-binding CD8 T cells (Figure 1C, Figure S1A-D). We observed abundant pMHC-binding to virus-related pMHC multimers, and less resolved recognition of NAg/CitNAgs (Figure 1D, Figure S1C-D) indicating that recognition of self-derived neuroantigens by CD8 T cells is mediated by low avidity interactions as shown previously in type 1 diabetes (Bulek et al., 2012). We furthermore observed a population of cross-binding CD8 T cells, driven in part by non-specific binding, as CD4 T cells showed HLA-I multimer cross-binding populations as well (Figure S1C-D). To avoid artefacts, we therefore focused our analysis on single multimer-positive CD8 T cells binding either virus-or NAg/CitNAg-derived epitopes going forward.

Epitopes from all antigens were recognized by CD8 T cells with varying estimated cellular frequency and prevalence (Figure 1E) with a high degree of epitope repertoire overlap between HD, RRMS and PMS groups for epitopes recognized by more than one donor (Figure S1E-F). The highest observed cellular frequencies of pMHC-binding CD8 T cells were CEF-specific CD8 T cells followed by laEBV-, NAg- and CitNAg-specific populations with median frequencies of 0.091%, 0.012%, 0.006% and 0.003%, respectively (Figure 1F). The rare cellular frequency of NAg and CitNAg-specific CD8 T cells coincided with an overall rarer prevalence of recognition per epitope, e.g., more than half of NAg- and CitNAg epitopes were recognized in single donors (Figure S1I). Consequentially, epitopes recognized in less than 5 donors made up more than 50% of the cumulative observed cellular frequency for NAg- and CitNAg-specific CD8 T cells (Figure 1G). Many laEBV-derived epitopes were also recognized in single donors (41 out of 87 epitopes) (Figure S1E), however, 50% of the cumulative observed cellular frequency was only reached when including responses observed in more than 15 donors (Figure 1G), suggesting that in contrast to NAg/CitNAg-specifc CD8 T cells, that the majority of laEBV-specific CD8 T cells discovered in peripheral blood recognized the same set of epitopes.

Overall, these data indicate that neuroantigen-specific CD8 T cells exist and recognize a diverse set of self-epitopes at low peripheral cellular frequency. We furthermore reason that antigen-driven expansion of immune dominant CD8 T cells recognizing epitopes from MAG, MBP, MOG, PLP, and TALDO1 is generally a rare event in peripheral blood compared to recognition of CEF and laEBV antigens.

### Lower cellular frequency of epitope-specific CD8 T cells in context of HLA-B*07:02 characterizes multiple sclerosis patients

Monitoring the cellular frequency of antigen-specific CD8 T cells in peripheral blood towards a wider panel of epitopes could reveal disease-specific patterns of NAg/CitNAg and virus antigen recognition in MS valuable for further longitudinal studies to substantiate the longitudinal data obtained previously (Angelini et al., 2013; Jilek et al., 2008; Pender et al., 2017).

The frequency of NAg-specific CD8 T cells obtained through flow cytometry was slightly reduced in patients with MS compared to healthy donors (Figure 2A). This was further supported by a lower median frequency of individual NAg-specific CD8 T-cell responses in RRMS and PMS patients compared to healthy donors, when measured by sequencing of multimer-associated DNA-barcodes (Figure 2B). A similar pattern was true for CitNAg-specific CD8 T cells, although only significantly for the PMS patients (Figure 2B).

**Figure 2:**
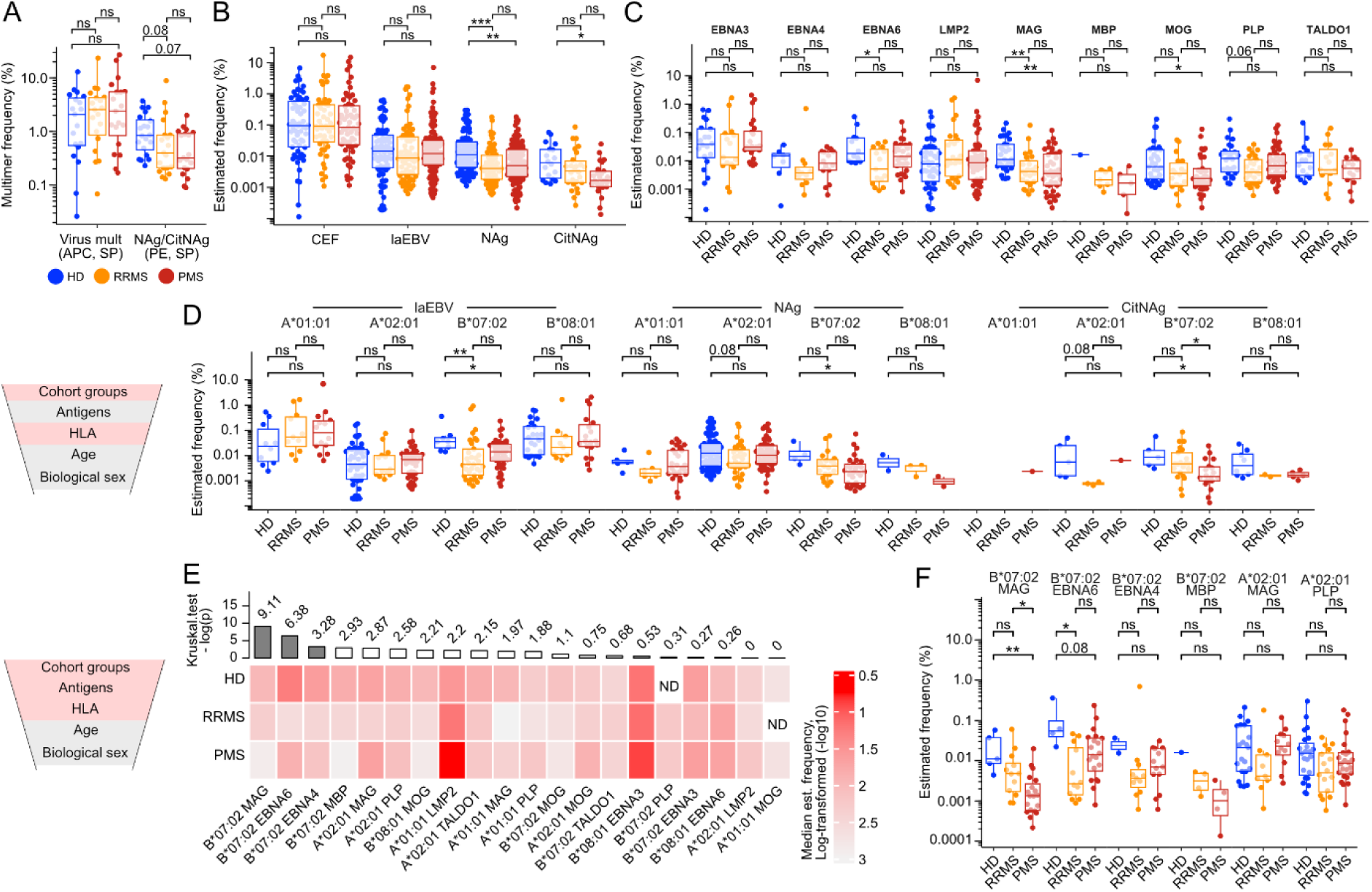
Differential cellular frequency of antigen-specific CD8 T cells in MS. A) Comparison of flow cytometry-based frequencies of single positive (SP) CD8 T cells binding either NAg/CitNAg-or virus-derived pMHC multimers among cohort groups. Color legend is representative for HD, RRMS and PMS patients throughout the figure. B) Comparison of read-based estimated frequencies of CD8 T cells binding to a given multimer for each group of antigens (CEF, laEBV, NAg and CitNAg) among cohort groups. Estimated frequency per multimer was calculated as described in materials and methods. C) Read-based estimated frequencies per antigen and cohort group. Both CitNAg- and NAg-specific responses were included for each autoantigen. D) Comparison between cohort groups of NGS-based estimated frequencies per HLA and antigen group. E) Heatmap comparing the median estimated frequency per HLA and antigen combination between cohort groups. Estimated frequencies among multimer-specific responses per group were normalized using feature-scaling placing all values per HLA and antigen combination between 0 and 1. The alpha value for significance was set at 0.001 or – log(p) = 3. ND, non-detected. F) Post-hoc analysis using raw estimated frequency values of the top 5 most significantly variable HLA and antigen combinations. P-values in E were calculated using Kruskal-Wallis test. Other P-values were calculated using Dunn’s test and adjusted for multiple hypothesis testing using the Benjamini Hochberg approach. *, **, and *** represent p-values lower than or equal to 0.05, 0.01, and 0.001 respectively. Near-significant adjusted p-values are indicated by numbers rounded to the nearest second digit. ns, non-significant.

On the contrary, we saw no global reduction in the estimated frequency of laEBV-specific CD8 T cells between multiple sclerosis patients and healthy donors (Figure 2B). However, when our data was stratified for individual antigens, we observed that EBNA6-specific CD8 T cells occurred at a lower estimated frequency in RRMS patients, specifically (Figure 2C). Furthermore, we find that MOG, PLP, and especially MAG-specific CD8 T cells contributed to the observed lower frequency of peripheral NAg-specific CD8 T cells in patients compared to healthy donors (Figure 2C). Observations stratified by antigen were relatively robust given that all antigens and cohort groups were including at least 5 independent donors apart from EBNA4 and MBP (Figure S2A) with relatively equal contributions among donors within each group (Figure S2B). On average, we observed 72.6 epitope-specific CD8 T-cell responses per antigen, although only 32 and 13 responses were recorded for EBNA4 or MBP, respectively (Figure S2C).

As multiple factors may contribute or confound our findings through dependency on antigen, HLA, age, duration disease and biological sex of the individual, we further stratified our findings by each of these parameters. We found that the largest frequency differences across CEF, laEBV, NAg, and CitNAg antigen-specific responses were observed in context of specific HLA alleles (Figure S2D) suggesting that any global effect between cohort groups is heavily dependent on HLA restriction.

We therefore stratified for both cohort group and HLA allele identities. We observed that the frequency of laEBV-specific CD8 T cells restricted to HLA-B*07:02 were specifically reduced in multiple sclerosis patients (Figure 2D), and that the frequency of NAg- and CitNAg-specific CD8 T cells in context of HLA-B*07:02 was reduced in PMS patients compared to healthy donors (Figure 2D). To explore the common contribution of HLA and antigen identity further, we additionally used the negative logarithmic p-value from a Kruskal-Wallis test to rank all possible frequency comparisons between combinations of HLA allele and antigen across HD, RRMS and PMS cohort groups with a conservative significance threshold of p = 0.01 (Figure 2E). Although the cohort size chosen for this study did not generally allow for additional stratification beyond one variable when comparing differences between cohort groups, we observed that MAG-specific CD8 T cells restricted to HLA-B*07:02 were significantly reduced in the periphery of patients with PMS, while CD8 T cells specific for EBNA6 were reduced in patients living with RRMS (Figure 2E-F). Although statistically non-significant after adjustment for multiple hypothesis testing, we also found a similar trend for B*07:02-restricted EBNA4-specific CD8 T cells in RRMS patients (HD vs RRMS, p = 0.04, p.adj = 0.131) (Figure 2F).

In summary, we find that the frequency of B*07:02-restricted CD8 T cells recognizing both NAg/CitNAg and laEBV epitopes from MAG and EBNA6, respectively, are depleted in patients living with MS in line with a previous report on B*07:02-restricted immune dysfunction in context of multiple sclerosis (Jilek et al., 2012).

### HLA shapes the antigen preference of CD8 T cells

Interestingly, for each HLA allele the observed frequency of laEBV- and NAg-specific responses, displayed a different, but yet discrete frequency ranges. HLA-A*02:01 showed the lowest median frequency of laEBV-specific CD8 T cells, while it simultaneously demonstrated the highest median frequency of NAg-specific CD8 T cells (Figure S2E). HLA genotypes furthermore skewed the antigen-preference of observed immune responses. For example, we observed that recognition of EBV in context of HLA-A*01:01 was largely restricted to LMP2 epitopes, whereas recognition in restriction to B*07:02 was limited to EBNA3, EBNA4, and EBNA6 (Figure S2F). This suggested that each antigen was, to a large degree, only recognized in context of a subset of the included HLA alleles. Indeed, EBNA3 was almost exclusively recognized in context of HLA-B*07:02 and B*08:01. EBNA4 was primarily recognized with HLA-B*07:02. EBNA6 was primarily recognized in context of HLA-B*07:02 and B*08:01. LMP2 was primarily recognized in context of HLA-A*01:01 and A*02:01. MAG was primarily recognized with HLA-A*02:01 and B*07:02. Furthermore, MBP was scarcely recognized with 7 out of 13 responses being towards citrullinated peptides observed in context of HLA-B*07:02 (all 13 peptides were of different amino acid sequences). MOG was primarily recognized in context of HLA-A*02:01 and B*07:02. PLP was recognized across HLA-A*01:01, A*02:01 and B*07:02, whereas native and citrullinated TALDO1-derived epitopes were recognized with A*02:01 and B*07:02, respectively (Figure S2G).

In summary, we find that HLA restriction strongly influences the frequency and target epitope repertoire of observed antigen-specific CD8 T cells.

### Interaction between age and antigen-specific cellular frequency in MS

As age had a non-negligible effect on the cellular frequency of NAg/CitNAg-specific CD8 T cells (Figure S2D), we further explored whether frequency of laEBV, NAg, and CitNAg-specific CD8 T cells was dependent on age or the duration of disease. Using the sum of estimated frequency as a measure of total antigen-specific CD8 T cells recognizing any given antigen, we find that the total frequency of NAg-specific and CitNAg-specific CD8 T cells weakly correlates with age (Figure S3A-B) and disease duration (Figure S3C-D), while laEBV-specific responses were independent of age and disease duration (Figure S3B, Figure S3D). We furthermore found a trend whereby the majority of cumulatively observed laEBV-specific responses were situated in younger healthy donors and older MS patients, respectively (Figure 3A-B), suggesting that young MS patients do not adequately control latent EBV virus infection, as previously suggested for newly diagnosed MS cases (Pender et al., 2017). Other minute cumulative patterns were also observed. The frequency of CMV-specific CD8 T cells rose steadily in healthy donors but lacked behind in RRMS and PMS patients (Figure 3A-B). Additionally, we observed that NAg-specific responses appeared to lack behind in PMS donors compared to both healthy donors and RRMS donors (Figure 3A). Interestingly, when the cumulative frequency was observed per antigen across different ages, we observed that EBNA6, LMP2, and MOG-specific CD8 T cells strongly contributed to the global discrepancy between MS patients and healthy donors (Figure 3B) indicating that disease-specific interactions with age may relate to, or correlate with, recognition of these antigens. In summary, our analysis circumstantially indicates a weak interaction between age and antigen-specific frequency of laEBV and NAg/CitNAg-specific immune responses, where people living with MS are characterized by a slower development of laEBV-specific immune responses compared to healthy donors.

**Figure 3:**
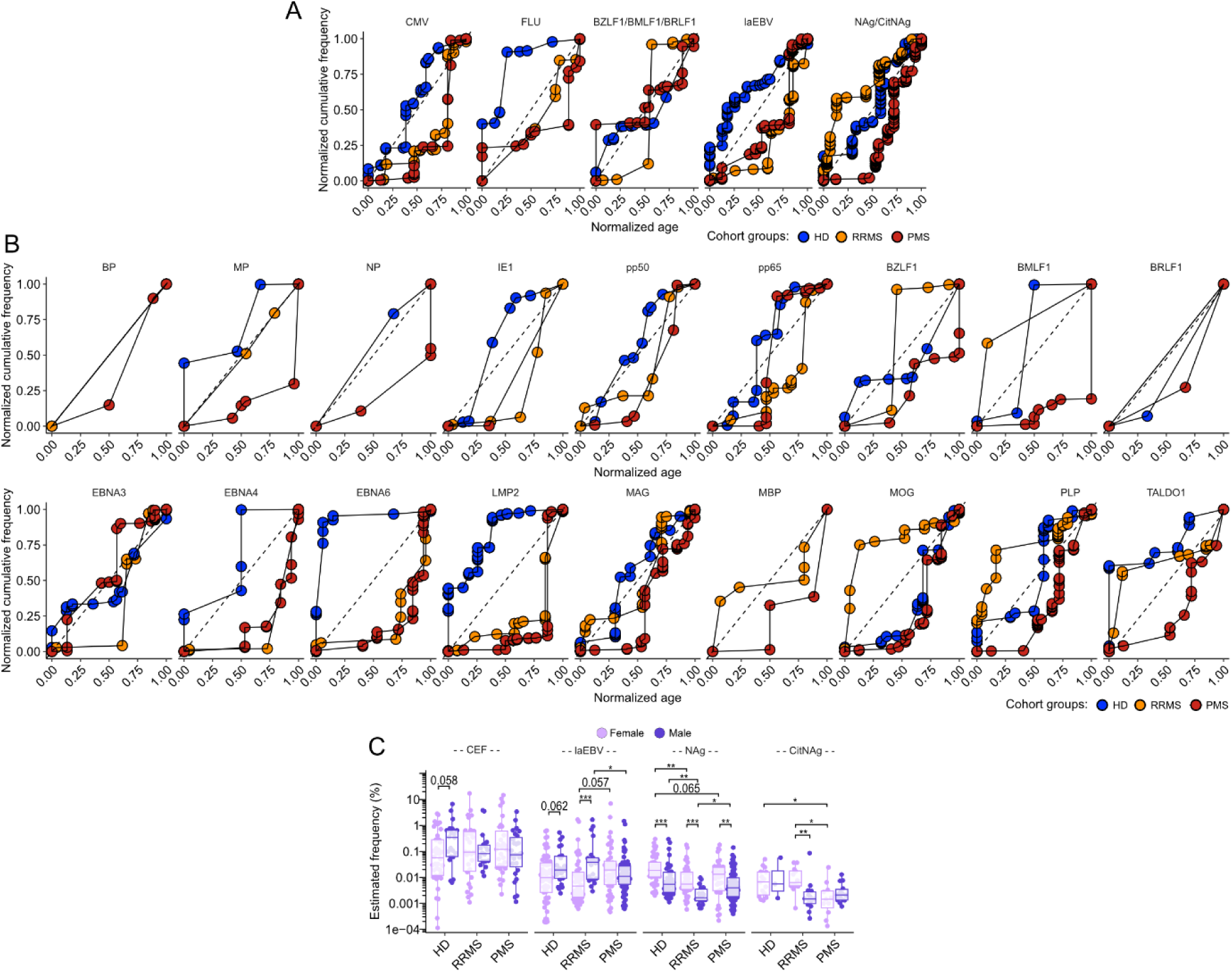
Age and biological sex as possible confounders of antigen-specific frequency in MS. A) Normalized cumulative frequency of antigen-specific CD8 T cells as a function of increasing group-wise normalized age (feature-scaling). B) As B but stratified per antigen. A single MBP-specific response amongst HD donors and a single NP-specific response in RRMS patients could not be visualized alone and was therefore excluded. C) Estimated frequency of all laEBV and NAg-specific immune responses stratified by biological sex and cohort groups. All unlabeled comparisons within and across cohort groups were statistically non-significant. P-values were calculated using a Dunn‘s test. *, **, and ***, represented p-values lower than 0.05, 0.01, and 0.001, respectively. Exact p-values were shown for near-significant comparisons.

### Interaction between biological sex and antigen-specific cellular frequency in MS

We also stratified our observations according to biological sex, which, like age, showed non-negligible effects in context of NAg-specific CD8 T-cell responses (Figure S2D). Female and male RRMS patients were characterized by a lower frequency of NAg-specific CD8 T cells compared to healthy females and healthy males, respectively, indicating that the RRMS-specific reduction of NAg-specific CD8 T cells is a general phenomenon to both sexes. We furthermore observed that female individuals were biased towards higher NAg-specific frequency among healthy donors and MS patient groups compared to their male counterparts (Figure 3C) suggesting that female individuals produce a higher number of NAg-specific CD8 T cells independent of disease. Interestingly, the observed NAg-specific frequency bias was only observed in context of HLA-A*02:01 and HLA-B*07:02 without noticeable bias in context of HLA-A*01:01 and B*08:01 (Figure S3E).

We also observed sex-specific differences among laEBV-specific CD8 T-cell responses, where especially female RRMS donors were characterized by reduced frequencies of laEBV-specific CD8 T cells compared to males with similar trends observed for female and male healthy donors (Figure 3D). However, sex-specific differences in laEBV-specific CD8 T-cell frequency were not observed among PMS donors (Figure 3D). This indicates that the frequency of laEBV-specific CD8 T cells to some extend depends on biological sex, and that female RRMS patients produce a lower number of laEBV-specific CD8 T cells compared to male RRMS patients. Like NAg-specific CD8 T cells, these statistical trends for laEBV-specific CD8 T cells were most pronounced in the context of HLA-A*02:01 and B*07:02 (Figure S3E).

Overall, our data indicates that biological sex skews NAg- and laEBV-specific immune responses towards a higher peripheral frequency of NAg-specific CD8 T cells in women, and a reduced frequency of laEBV-specific CD8 T cells in context of HLA-A*02:01 and B*07:02.

### Patients living with MS largely recognize the same set of epitopes as healthy individuals with important exceptions

Patients living with MS could hypothetically recognize a unique epitope, unrecognized in healthy donors, and vice versa. We therefore estimated the group-wise prevalence of recognition of individual CD8 T-cell epitopes, i.e. the fraction of donors with positive epitope-recognition for each of the cohort groups (HD, RRMS and PMS) (Figure 4A), followed by a Fisher’s exact test to annotate significant prevalence of recognition in three pairwise analyses (Figure 4B-D, supplemental table 3).

**Figure 4:**
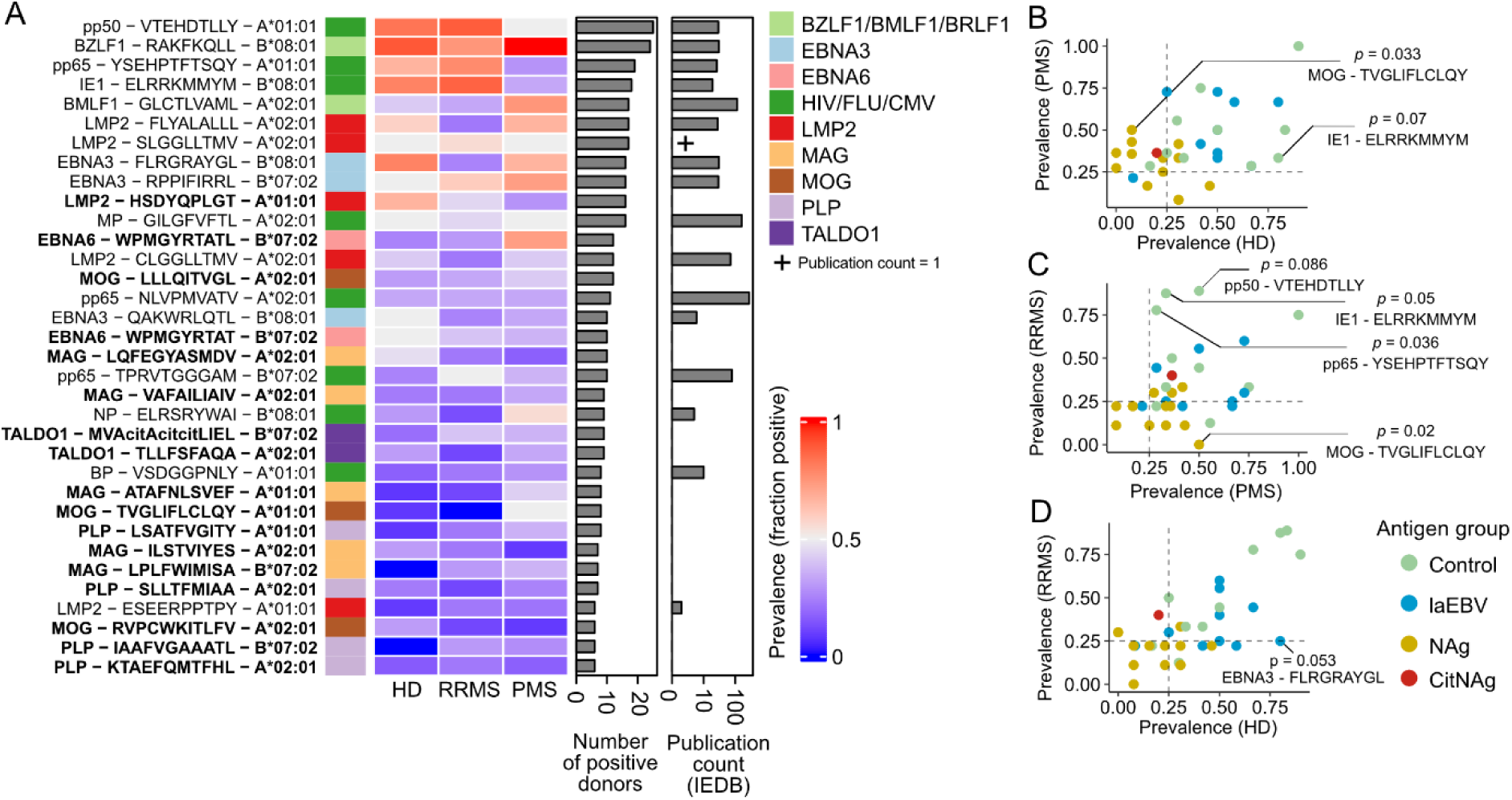
Commonly recognized epitopes by healthy donors and MS patients. A) A heatmap showing the observed prevalence of recognition of each epitope recognized in more than 5 independent individuals (n = 34 epitopes). Columns from left to right show antigen of origin, prevalence in healthy donors, prevalence in RRMS patients, prevalence in PMS patients, number of positive donors, and publication count according to IEDB, respectively. Prevalence of recognition is reported as a positive fraction of tested individuals. Bold text indicates novel epitopes unreported in IEDB. B) Differential prevalence of recognition between RRMS patients and healthy donors. C) Differential prevalence of recognition between PMS patients and healthy donors. C) Differential prevalence of recognition between RRMS patients and PMS patients. Only epitopes from the minimal epitope panel in A are included in B-D. P-values were calculated using an unadjusted Fischer‘s exact test to indicate statistical trends.

Multimer-binding CD8 T cells were prevalently detected for BZLF1-RAKFKQLL and pp50-VTEHDTLLY with CD8 T-cell responses observed in more than 20 donors (Figure 4A). However, whereas recognition of BZLF1-RAKFKQLL was observed in almost all HLA-matched donors (HD: 90%, RRMS: 75%, PMS: 100%) recognition of pp50-VTEHDTLLY (HD: 83.3%, RRMS: 88.9%, PMS: 50%) and other CMV-derived epitopes pp65-YSEHPTFTSQY (HD: 66.7%, RRMS: 77.8%, PMS: 53.2%) and IE1-ELRRKMMYM (HD: 80%, RRMS: 87.5%, PMS: 33.3%) trended towards lower recognition prevalence in PMS patients compared to both RRMS and HD donors (Figure 4B-D).

We furthermore observed that MOG-TVGLIFLCLQY (HD: 7.69%, RRMS: 0%, PMS: 50%) and EBNA3-FLRGRAYGL (HD: 80%, RRMS: 25%, PMS: 66.7%) exhibited differential prevalence of recognition among our cohort groups (Figure 4A-D). Recognition of MOG-TVGLIFLCLQY restricted to HLA-A*01:01 was rare in healthy donors (1 out of 13 donors), absent in RRMS donors (0 out of 9 donors) yet recognized in 50% of HLA-matched PMS patients (7 out of 14 donors) indicating a specific enrichment of MOG-TVGLIFLCLQY responses in PMS donors. EBNA3-FLRGRAYGL restricted to HLA-B*08:01 was recognized by 80% of healthy donors (8 out of 10 donors), 25% of RRMS donors (2 out of 8 donors), and 66.7% of PMS donors (6 out of 9 donors) indicating a specific absence of EBNA3-FLRGRAYGL responses from RRMS donors.

Note that BZLF1-RAKFKQLL and EBNA3-FLRGRAYGL represent two common epitopes recognized by cytotoxic CD8 T cells restricted to HLA-B*08:01 from the lytic and latent cycles of EBV infection, respectively. These were previously reported in a patient refusing disease-modifying therapy, where active disease was associated with a relative expansion of peripheral antigen-specific CD8 T cells targeting BZLF1-RAKFKQLL and a loss of EBNA3-FLRGRAYGL-specific CD8 T cells over time (Angelini et al., 2013).

Finally, we were interested in recording any novel epitopes previously undescribed in the literature. To this end, we downloaded and integrated publication record data per epitope reported within the Immune Epitope database (IEDB)(Vita et al., 2019). We found several unreported epitopes that were prevalently recognized in our study while absent in IEDB. These included HLA-A*01:01-restricted LMP2-HSDYQPLGT (HD: 66.7%, RRMS: 44.4%, PMS: 28.6%), HLA-B*07:02-restricted EBNA6−WPMGYRTATL (HD: 25%, RRMS: 30%, PMS: 72.7%) and EBNA6−WPMGYRTAT (HD: 50%, RRMS: 40%, PMS: 36.4%) - a length variant of EBNA6−WPMGYRTATL (Figure 4A). Additionally, many of the most prevalently recognized epitopes such as BZLF1-RAKFKQLL and pp50-VTEHDTLLY were also well reported within the IEDB with more than 20 publication records in the past.

Taken together, we described the prevalence of recognition for both common as well as novel epitopes and find that healthy donors, as well as people living with MS, largely recognize the same set of laEBV and NAg/CitNAg epitopes with a tendency for RRMS patients have undetectable for EBNA3-FLRGRAYGL recognition, and for PMS to recognize fewer CMV-related epitopes than healthy. Finally, MOG-TVGLIFLCLQY represented an interesting outlier specifically recognized by PMS patients.

### Neuroantigen-specific CD8 T cells are enriched for anergic, naïve, and long-lived central memory phenotypes

Phenotypes of antigen-specific CD8 T cells occur in humans on a scale from anergic, naive and less differentiated memory phenotypes to senescent or terminally differentiated phenotypes with or without effector functions (Galluzzi et al., 2025). To observe the differentiation status of virus- and neuroantigen-specific CD8 T cells observed here, we acquired immunophenotyping data for 10 healthy donors, 9 RRMS, and 10 PMS patients in addition to staining with pMHC multimer libraries. In our assay, NAg/CitNAg-specific CD8 T cells exhibited a ∼2.2-fold lower surface expression of the CD8 co-receptor compared to both the multimer-negative background and virus-specific CD8 T cells (Figure 5A-B). Additionally, NAg/CitNAg-specific CD8 T cells showed higher expression of CD127 (IL7Rα) (Figure 5C). Notably, both virus-specific and NAg/CitNAg-specific CD8 T cells displayed similar surface expression of CD8 and CD127 expression in healthy donors and individuals with MS (Figure S4A-B).

**Figure 5:**
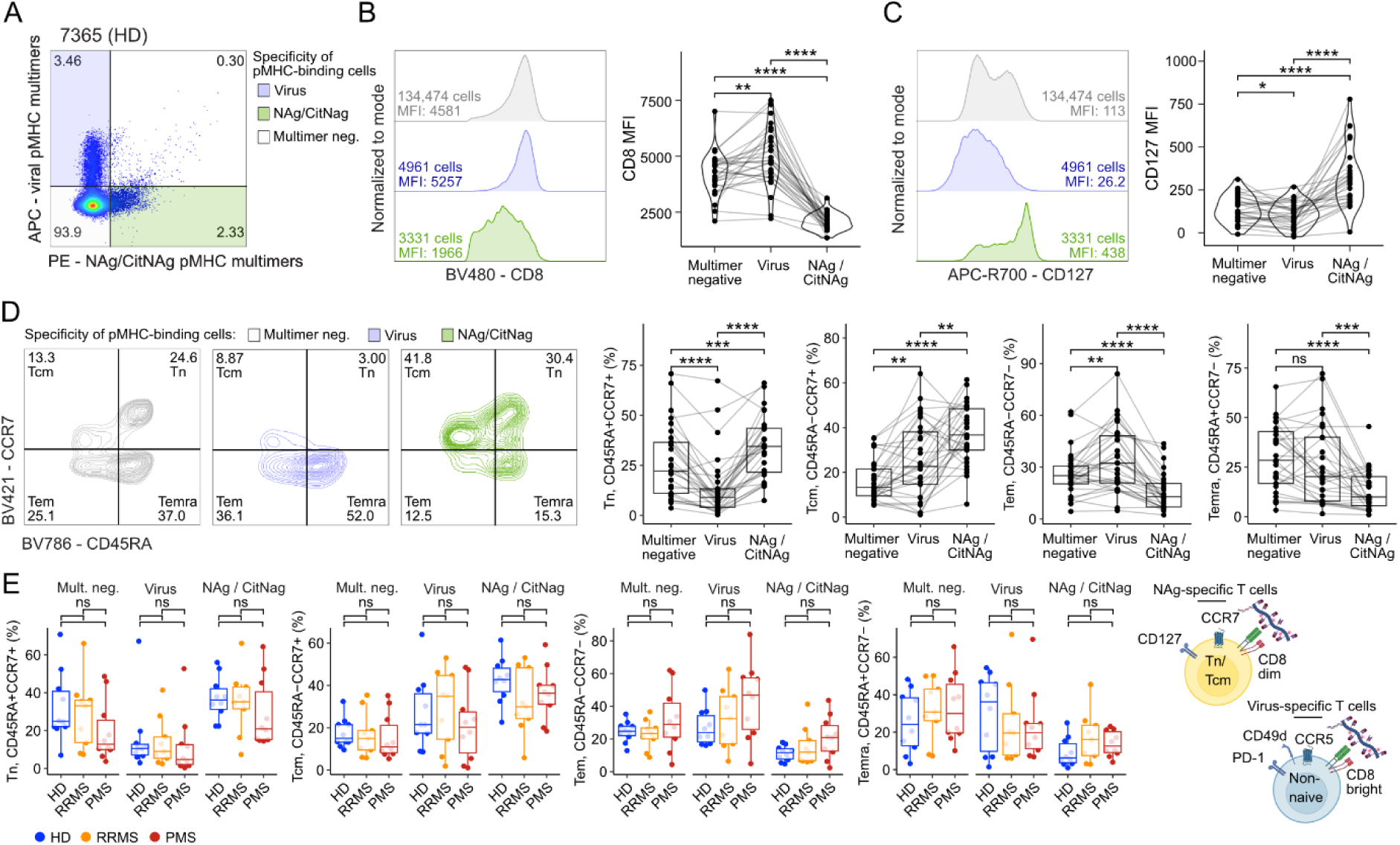
Immunophenotypes associated with virus- and NAg-specific CD8 T cells in MS patients and healthy donors. A) Example plot of pregating pertaining to virus and NAg/CitNAg-specific CD8 T cells. B) Representative histogram and paired quantifications of CD8 surface expression using median fluroescence intensity (MFI). C) Representative histogram and paired quantifications of CD127 surface expression using MFI. D) Representative contour plots and paired quantifications of CCR7+CD45RA+ differentiation stages Tn, Tcm, Tem, Temra as indicated. E) Comparison of differentiation stage frequencies among cohort groups (HD, RRMS, PMS) across multimer negative (mult. neg) reference CD8 T cells, virus-specific CD8 T cells (Virus), and NAg/CitNAg-specific CD8 T cells (NAg/CitNAg). Each dot represents a single donor. P-values were calculated using a pairwise Wilcox test in B-D, while a Dunn’s test was used for E. All p-values were adjusted using the Benjamini-Hochberg approach. *, **, ***, and **** represented p-values lower than 0.05, 0.01, 0.001, and 0.0001, respectively. ns, non-significant. Biorender illustrations were created in BioRender Hadrup, S. (2025) https://www.biorender.com/zzbgokl.

In contrast to NAg/CitNAg-specific CD8 T cells, we find that virus-specific CD8 T cells exhibited increased surface expression of the CD8 co-receptor and decreased expression of CD127 compared to multimer-negative cells, which is in line with these cells being enriched for effector-memory subsets (Tem, CCR7-, CD45RA-) (Figure 5D-E). NAg/CitNAg-specific CD8 T cells were generally non-differentiated compared to virus-specific and multimer-negative cells with the majority of cells exhibiting naive-like (Tn, CCR7+,CD45RA+) or central-memory phenotypes (Tcm, CCR7+, CD45RA-) while effector memory subsets including effector memory T cells re-expressing CD45RA (Temra, CCR7-, CD45RA+) were low frequent (Figure 5D-E). The high enrichment of CCR7+ subsets among NAg/CitNAg-specific CD8 T cells would suggest that these are circulatory naive-like cells, which is further supported by their low surface expression of integrin α-4 (CD49d) (Figure S4C-D). Finally, NAg/CitNAg-specific CD8 T cells expressed slightly lower level of PD-1 and higher level of CD20 compared to multimer-negative populations, while maintaining the same amount of CD57 and CCR5 (Figure S4E-K), whereas virus-specific CD8 T cells expressed significantly higher amounts of CD49d, PD-1, CD20 and CCR5 compared to their multimer negative counter parts (Figure S4E-K).

Taken together this suggests that NAg/CitNAg-specific CD8 T cells exhibit a circulatory, naive-like state in addition to an anergic phenotype with decreased sensitivity to antigen-stimulation previously associated with a decreased response to antigen-stimulation marked by a reciprocal downregulation of CD8 and upregulation of CD127 expression are associated with a state of anergy to antigen-stimulation (Park et al., 2007).

Importantly, we do not find any phenotypic differences between antigen-specific CD8 T cells found MS patients compared to healthy donors in regard to differentiation status, i.e. CCR7 and CD45RA expression (Figure 5E), CD49d surface expression (Figure S4D), PD1 surface expression (Figure S4F), CD57 surface expression (Figure S4H), or CD20 expression (Figure S4J). That said, we find a trend for both multimer-negative and multimer-positive cells to express more CCR5 in PMS patients compared to healthy donors (Figure S4L).

### Verification of low-frequent antigen-specific CD8 T cells using peptide-based in vitro expansion

To verify the presence of low-frequent antigen-specific CD8 T cells recognizing our identified peptide epitopes we stimulated and expanded antigen-specific CD8 T cells from peripheral blood from 3 MS patients over 14 days using peptide stimulation of PBMCs. Specifically, we produced unstimulated, or peptide-pool stimulated in vitro expansion cultures, where T cells were either stimulated with a laEBV-specific peptide pool or a NAg/CitNAg-specific peptide pool testing for expansion of responses associated with HLA-A*02:01, B*07:02 and B*08:01. After 14 days of in vitro expansion, we used pMHC multimers to enumerate epitope-specific responses.

Virus-specific CD8 T cells clearly harbored high expansion potential after cognate peptide stimulation compared to unstimulated control cultures (Figure 6A), whereas NAg/CitNAg-specific immune responses largely remained low frequent or absent from the culture following expansion (Figure 6B). Nevertheless, by looking at individual pMHC multimer stains (Figure 6C, Figure S5), we verified two PLP-derived NAg-restricted immune responses restricted to HLA-A*02:01 PLP-FMIAATYNFAV and PLP-YALTVVWL, as well as A*02:01-restricted MAG-VAFAILIAIV (Figure 6C) – where the latter expanded from ∼0.003% to ∼0.03% (10-fold) after stimulation with virus peptides hinting at cross-reactivity, although we could not identify the partner epitope in the pool of virus epitopes we used. The absence of multimer+ cells after expansion may have to do with their naive-like / anergic phenotype, which likely prevented us from validation of additional NAg-specific CD8 T-cell populations.

**Figure 6:**
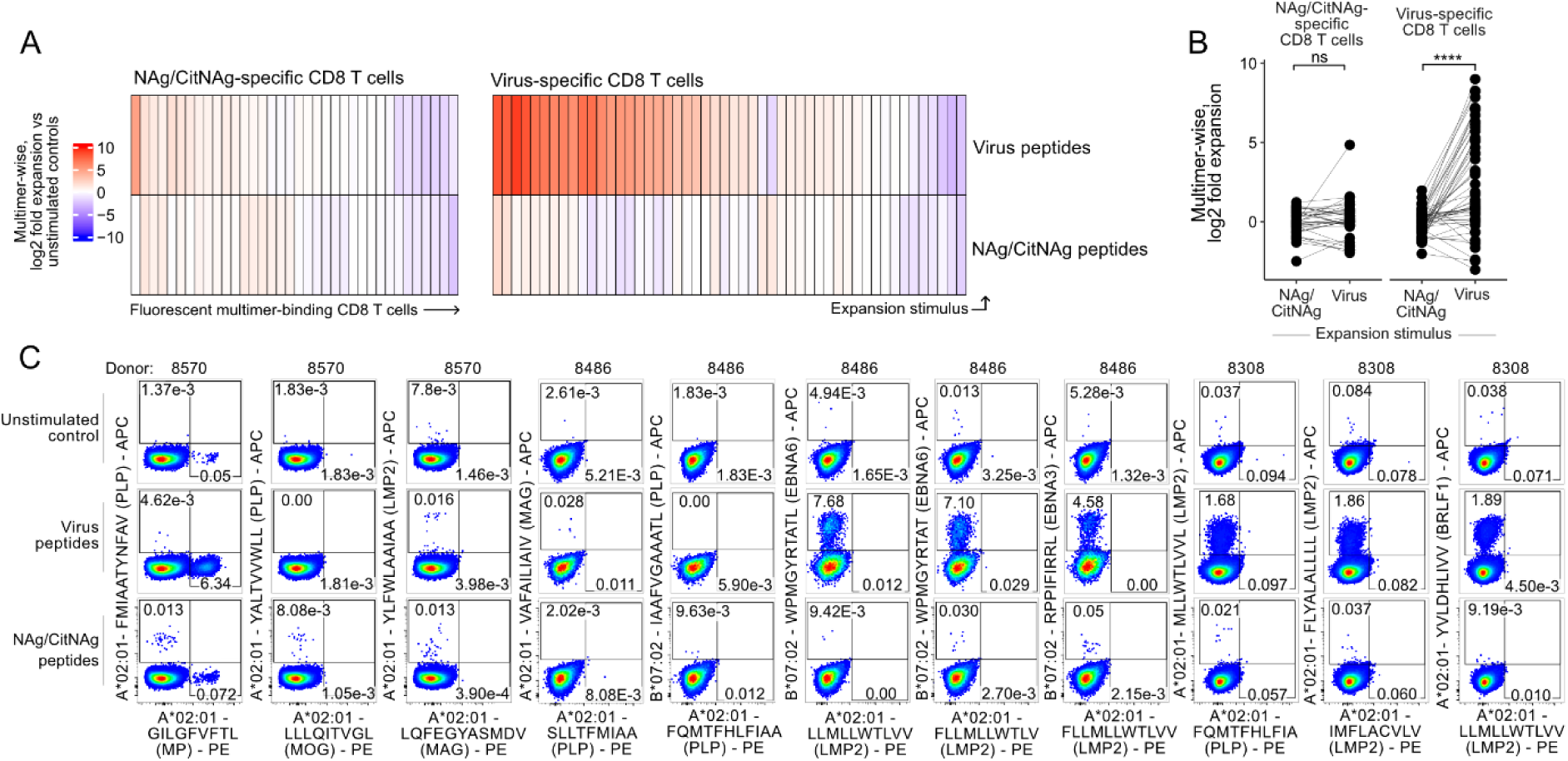
NAg-specific CD8 T cells show limited proliferation in response to peptide stimulation. A) Log2-fold expansion as measured by dextran pMHC multimer staining after 14 days of expansion using peptide pools of select virus and NAg-derived peptides detected in one or more donors. Multimer-wise expansion is calculated as a log2 fold change from the unstimulated baseline. B) Group-wise comparison of expansion comparing the effect of using virus peptides or NAg/CitNAg-derived peptides as expansion stimulus. C) Select populations of antigen-specific CD8 T cells following peptide-based expansion showing notable differences between expansion conditions. P-values in B were calculated using a paired Wilcox test. **** represented a p-value lower than 0.0001. ns, non-significant.

Strikingly B*07:02-restricted EBNA6-WPMGYRTATL and EBNA6-WPMGYRTAT-specific CD8 T cells both expanded heavily over the course of our 14-day expansion from ∼0.004% in unstimulated cultures to ∼7.4% after cognate virus peptide stimulation (Figure 6C). In comparison, EBNA3-RPPIFIRRL-specific CD8 T cells of the same donor expanded from ∼0.005 to ∼4.6% following virus peptide stimulation. As EBNA3-RPPIFIRRL is a commonly recognized EBV peptide restricted to HLA-B*07:02 (Kristensen et al., 2024), our EBNA6-WPMGYRTAT(L) peptides likely constitute a similarly common epitope from EBV infection recognized in context of HLA-B*07:02. Finally, we also observed a sizeable population of LMP2-MLLWTLVVL-specific CD8 T cells after expansion, which represented an epitope unreported in IEDB and recognized in less than 5 independent HLA-A*02:01-positive donors (Figure 6C).

Taken together, we verified multiple novel epitopes in context of HLA-A*02:01 and B*07:02 recognized by antigen-specific CD8 T cells with high expansion capacity while underscoring a tendency for NAg/CitNAg-specific CD8 T cells to not respond to antigen-stimulation in vitro.

## DISCUSSION

This study presents a detailed map of both laEBV and NAg-derived T-cell epitopes, of relevance for further longitudinal studies of antigen-specific CD8 T cells in MS and beyond. We found that T cells in MS patients and healthy donors largely recognize the same set laEBV and NAg-derived epitopes except for in the context of HLA-B*07:02. Here, recognition of laEBV EBNA6 and NAg MAG was specifically reduced in patients living with MS. This matches previous observations, where HLA-B*07:02 drives a weak form of EBV-specific hypocytosis in MS patients (Jilek et al., 2012).

Our observations also provide a nuance to the data reported by Pender et al (Pender et al., 2017) as we found that the global frequency of laEBV-specific CD8 T cells in MS patients was overall similar to healthy donors. Pender and colleagues used peptide pools of latent and lytic antigens and PBMC stimulation to enumerate %IFNγ antigen-reactive CD8 T cells and found a skewing of the immune response in MS patients towards higher frequencies of latent antigen-reactivity in MS patients and a lower frequency of lytic-specific CD8 T cells in MS patients compared to healthy donors (Pender et al., 2017). However, similar to our data, they only found slight differences when using pMHC multimers to enumerate laEBV-specific CD8 T cells. Suggesting that differences in PBMCs are minor and possibly related to functional capabilities, rather than actual T-cell frequencies.

To further investigate the hypothesis that MS disease manifestation and relapse is a consequence of dysfunctional EBV control, further immunomonitoring studies in humans could focus on using epitopes discovered herein supplemented by epitopes discovered from lytic phase EBV antigens as well as epitopes restricted to HLA class II and to HLA-E to longitudinally monitor EBV-specific CD8 T-cell frequency and function in MS in a multiomic single-cell setting.

This would be especially interesting in the two specific longitudinal settings as reported by Angelini and colleagues (Angelini et al., 2013). In one setting, two patients refused treatment but permitted longitudinal assessment of EBV-specific T cells throughout disease progression. Here, lytic antigen-specific CD8 T cells proliferated and subsequently contracted within months of active disease. In another setting, patients permitted immunomonitoring of antigen-specific CD8 T cells before and after initiation of disease-modifying therapy (Natalizumab) showing a stark increase in laEBV-specific CD8 T cells after initiating therapy (Angelini et al., 2013), which may in part be due to the retention of tissue-homing memory T cells from the CNS compartment. Given our data as reported here, important epitopes to include in future immunomonitoring is EBNA6-WPMGYRTAT(L) and EBNA3-FLRGRAYGL both showing an RRMS-specific reduction in specific CD8 T cells compared to healthy controls; a reduction similar to the patient-specific reduction in the CNS-homing subset of T9 CD4 T cells compared to healthy controls identified by Kaufmann and colleagues (Kaufmann et al., 2021).

Interestingly, HLA-B*07:02 is frequently associated with the HLA-DRB1*15:01 through the prevalent A3-Cw7-B7-DR15-DQ6 haplotype (Chao et al., 2007). As only a single healthy donor was positive for HLA-DRB1*15:01, we can therefore not exclude the possibility of HLA-DRB1*15:01 exerting an effect on the frequency of antigen-specific CD8 T cells observed here. That an MHC class II risk allele would exert an effect on the observed frequency of antigen-specific CD8 T cells has been observed previously by us in context of narcolepsy and HLA-DQB1*06:02 (Pedersen et al., 2019) as well as by others in humanized animal models (Zdimerova et al., 2021), and so we cannot rule out that the observed differences between B*07:02+ healthy donors and B*07:02+ MS patients is due to the presence and absence of HLA-DRB1*15:01 in patients and healthy donors, respectively.

Broadly screening for recognition of candidate epitopes in a highly multiplexed setting can allow for the definition of unique epitopes recognized by MS patients and unrecognized in healthy donors. Here, we largely observed a similar landscape of epitope recognition with prevalent recognition of the same set of epitopes, however, MOG-TVGLIFLCLQY recognized in context of HLA-A*01:01 was specifically enriched in patients living with PMS – the exact group with the lowest overall frequency of NAg/CitNAg-specific CD8 T cells. The implications of this remain unclear. Moreover, we observed a statistical trend for more prevalent recognition of multiple CMV-derived epitopes including IE1-ELRRKMMYM, pp65-YSEHPTFTSQY, and pp50-VTEHDTLLY negatively associated with RRMS yet recognized in PMS and healthy donors supporting a protective indication by CMV-specific lymphocytes observed previously (Bjornevik et al., 2022; Vietzen et al., 2023)

We attempted to verify as many T-cell responses as possible in three donors through peptide-driven expansion of antigen-specific CD8 T cells. Through this, we verified the presence of LMP2-MLLWTLVVL-specific CD8 T cells restricted to HLA-A*02:01 in addition to the B*07:02-restricted EBNA6-WPMGYRTAT(L) epitopes. These constitute novel epitopes in the literature according to IEDB (Vita et al., 2019), although LMP2-MLLWTLVVL could be recognized by the same CD8 T cells recognizing LMP2-LLWTLVVLL, which according to IEDB is a much more frequently studied epitope.

A limitation to our study is the lack of specific verification of many NAg-specific CD8 T cells in addition to our highly multiplexed pMHC multimer assay. DNA-barcoded pMHC multimers are highly sensitive (Bentzen et al., 2016; Immudex, 2019; Ma et al., 2021), and it may well be that many NAg/CitNAg-specific CD8 T cells were unable to undergo peptide-based expansion due to their anergic phenotype (CD8dim CD127hi) (Park et al., 2007). The anergic phenotype is furthermore described by Maeda et al, where CD8 T cells capable of recognizing a HLA-A*02:01-restricted epitope derived from Melan-A is controlled by T regulatory cells (Tregs) while maintaining a naive-like (CCR7+, CD45RA+) phenotype (Maeda et al., 2014). NAg/CitNAg-specific CD8 T cells may similarly be controlled by Tregs although this remains speculative.

CD8 T cells including EBV-specific CD8 T cells have previously been found to survey the CNS (van Nierop et al., 2017; Serafini et al., 2019), with higher cell numbers observed during active inflammation in MS (Fransen et al., 2020). It is therefore relevant to speculate, which of the multimer-positive populations we describe with immunophenotyping, could infiltrate the CNS. NAg/CitNAg-specific CD8 T cells were enriched for CD49d-, CCR5dim and CCR7+ phenotypes, which strongly indicated that these are dominated by circulatory cells that are likely unable to reach the inflammatory conditions of the CNS without unspecific trauma or blood-brain barrier breakdown. That said, NAg/CitNAg-specific CD8 T cells showed a trend for higher CD20 surface expression, which has recently been associated with CNS migration especially in early MS (von Essen et al., 2025).

Virus-specific CD8 T cells showed relatively higher expression of CD49d, CCR5 and a lower expression of CCR7 compared to NAg/CitNAg-specific CD8 T cells and may therefore constitute the majority of antigen-specific CD8 T cells capable of tissue extravasation. Nevertheless, this does not exclude the possibility for NAg/CitNag-specific CD8 T cells to be primed in lymphoid organs associated with CNS drainage, e.g., the deep cervical lymph nodes. In general, analyses of tissues closer to the disease area is preferable, and the lack of a strong disease signature in PBMCs may simply be a consequence of location, as previously explored related to CD8 T-cell recognition in type-1-diabetes (Culina et al., 2018).

In conclusion, we report novel tendencies for a specific reduction in antigen-recognition of laEBV and NAg/CitNAg antigens by multiple sclerosis patients compared to healthy donors, which was especially enriched in context of HLA-B*07:02 along with a novel description of antigen-specific immunophenotypes associated with recognition of NAg/CitNAg-specific CD8 T cells.

## MATERIALS AND METHODS

### Patient samples and ethical approval

Density centrifugation was used to isolate PBMCs from peripheral blood samples from 20 healthy donors, 19 patients with RRMS, and 19 patients with PMS at the Danish Multiple Sclerosis Center at Copenhagen University Hospital – Rigshospitalet. Cells were cryopreserved in fetal calf serum (FCS) plus 10% dimethyl sulfoxide (DMSO). A summary of sex, age, duration of disease, EDSS scores and disease course of PMS patients is included in table 1. All patients were either treatment naive or had not received high-dose steroid treatment for at least one month and had not received immunomodulatory treatment for at least three months. None of the patients had ever received immunosuppressive drugs, e.g., cyclophosphamide, cladribine or mitoxantrone, or cell depleting monoclonal antibodies. All participants gave informed, written consent to participation. The study was approved by the regional scientific ethics committee (protocol number KF-01114309).

### Prediction of HLA binding peptides

Protein and isotype sequences were obtained from EBNA3 (P12977), EBNA4 (P03203), EBNA6 (P03204), LMP2 (LMP2A: P13285-1, LMP2B: P13285-2), MAG (P20916-1 and 2 isoforms), MBP (P02686 and 5 isoforms), MOG (Q16653 and 13 isoforms), PLP (P60201-1 and -2), TALDO1 (P37837) protein- and isoform sequences were obtained via https://Uniprot.org and used as input for netMHCcons1.1 (Karosiene et al., 2012). As inclusion criteria all peptides needed to have weak to strong predicted peptide affinities of EL_Rank 1 or lower. 2 umol lysophilized powder of each peptide was acquired through Pepscan/Biosynth (Netherlands), and resuspended in fresh DMSO and stored at -21°C. HLA class I-binding peptides were predicted for citrullinated neuroantigens (CitNAgs) by replacing each arginine with “X” in the given antigen to represent any amino acid moiety.

### Assembly of DNA-barcode labelled pMHC multimers

25mer DNA-barcodes designed as previously described (Bentzen et al., 2016; Xu et al., 2009) were acquired from LGC Biosearch Technologies (Denmark). In brief, two partially complementary single-stranded oligos (26 μM 5’Biotinylated C6-linker Ax oligo and 52 μM By oligo) were mixed with 5× Sequenase Reaction Buffer mix (70702, Thermo Fisher) and annealed at 65°C and slowly cooled to <35°C over 15-30 minutes followed by elongation over 10 minutes using Sequenase polymerase (70775Y, Thermo Fisher), 6.4 mM DTT (Thermo Fisher, R0861) and 1.28 mM dNTP (18427-088, Life Technologies) at room temperature. Elongated AxBy barcodes were diluted to 2.17 µM in nuclease-free water + 0.1% Tween and stored at -21°C in several alliquotes. Identities of Ax and By oligoes are available in Kristensen et al. (Kristensen et al., 2024).

APC-conjugated and PE-conjugated SA-Dextran multimers at approximately 500 kDa were custom made from Fina Biosolutions (USA) with the following sets of molecular ratios MOS596A: 2.95 APC/Dex, 3.9 SA/Dex and MOS722: 5.4 PE/Dex, 6.2 SA/Dex. Dextrans were diluted to 160 nM in PBS + 0.02% NaN3 before storage at 4°C and usage in multimer assembly. Dextrans were preincubated with biotinylated and elongated DNA-barcodes at 4°C for 30 minutes at stoichiometry of 0.5 DNA-barcodes per dextran. Exchange of UV-cleavable A*01:01, B*07:02, B*08:01 were then performed according to previously published protocols (Rodenko et al., 2006). Specifically, UV-cleavable monomers were exchanged at 50 ug/ml monomer and 100 µM peptide for 1 hr under 366 nm UV light. Cystein-stabilized A*02:01-Y84C were produced as described previously (Saini et al., 2019) and loaded by incubating monomers and peptides of interest at 50 ug/ml and 100 µM peptide respectively for 30 minutes at room temperature. Loaded biotinylated pMHC monomers were spun at 3300g, 4°C, for 5 minutes to sediments aggregates before addition to dextrans at 16 pMHCs per dextran. After incubation with pMHCs for 30 minutes, freezing buffer containing D-biotin, Herring DNA, EDTA, BSA, Glycerol and PBS to a final concentration among multimers of 5% glycerol, 1.5 µM D-Biotin, 0.5% BSA, 100 ug/ml herring DNA, and 2 mM EDTA. We routinely assembled 45 stains per batch of multimers. Each batch of multimers were tested individually for capacity to detect the same immunodominant CEF epitopes before using the multimer libraries to screen patient material. Virus-derived peptides were assembled on APC-SA-dextrans whereas self-derived peptides were assembled on PE-SA-dextrans.

### Staining with pools of barcode-labelled pMHC multimers and antibodies

Multimer libraries were thawed on ice and spun at 3300g, 4°C, 5 minutes. Supernatants were pooled in HLA-specific pools using 1.5 µL per stain, and split/recombined into donors-specific pools of HLA-matched multimer pools. These pools were then concentrated using vivo6 spin columns (VS0642, Sartorius, 100000 MWCO) and resuspended in barcode cytometry buffer (0.5% BSA, 100 ug/ml herring DNA, 2 mM EDTA) to 60 µL per stain + 10 µL. Multimer pools were spun 10.000 g for 2 min at 4°C in new 1.5 ml Eppendorf tubes, twice, followed by volume adjustment back to 60 µL per stain + 10 µL. A small aliquot of 5 µL was directly frozen at -21°C and used as reference for the barcode distribution in the original pool of multimers. Barcode-labelled multimers were then stored approximately 1 hour on ice before used for staining. PBMCs were thawed in parallel with multimer pooling using preheated RPMI 1640 (Gibco, 61870-044, Glutamax® supplemented) supplemented with 5% FCS (Gibco, 10500-064) and washed twice before resuspension in 96 well plates RPMI+5%FCS. Before addition of multimer pools to wells containing 5-10 million PBMCs, cells were washed with barcode cytometry buffer. All washes were performed at 390 g, 4°C, 5 minutes. Pellets were then resuspended using 60 µL multimer pools and 5 µL of 1 µM Dasatinib followed by brief incubation for 15 minutes at 37°C. Antibody mastermix were prepared in Brilliant Staining Buffer Plus (BD, 566385) subsequently added to each well. Two antibody master mixes were used throughout. Master mix #1 contained anti-human CD3-Pacific Blue (317313, Biolegend, OKT3), CD8-BV480 (566121, BD, RPA-T8), and LIVE/DEAD™ Fixable Near-IR (L34976, Invitrogen). Master mix #2 contained CD3-BUV395 (563546, BD, UCHT1), CD4-BUV563 (612913, BD, SK3), PD1-BUV737 (565299, BD, EH12.1), CRR7-BV421 (353233, Biolegend, G043H7), CD8-BV480 (566121, BD, RPA-T8), CD49d-BV605 (304323, Biolegend, 8F10), CCR5-BV650 (562576, BD, 2d7/CCR5), CD25-BV711 (302636, Biolegend, BC96), CD45RA-BV786 (563870, BD, HI100), CD20-PE-Cy5 (561761 BD,2H7), CD57-PE-Cy7 (393310, Biolegend, QA17A04), CD127-APC-R700(565185, BD, HIL-7R-M21) and LIVE/DEAD™ Fixable Near-IR (L34976, Invitrogen). Staining with surface antibodies was performed in 100-150 µl total volume. 34 donors were stained with both surface and intracellular anti-human FoxP3-PE-CF594 (562421, BD, 259D/C7) to which Invitrogen’s FoxP3 staining kit for intracellular staining (00-5523-00, Invitrogen) was applied following manufacturer’s protocol. Data from intracellular staining corresponded to data with surface antibodies confirming the presence of immunodominant CEF epitopes in both using material from technical controls. Fixed, permeabilized and stained cells were subsequently washed with barcode-cytometry buffer and stored at 4°C until sorting.

### Cell sorting and library preparation

BD Aria Fusion II was used to sort APC- and –PE multimer-binding CD3+CD4-CD8+ T cells. Single positive APC- and PE-positive CD8 T cells were sorted into the same Eppendorf tube containing 50 µL barcode cytometry buffer. Double positive cells were sorted into separate Eppendorf tubes. Sorted cells were subsequently pelleted by centrifugation at 5000g, 10 min, 4°C, and supernatant completely removed before freezing at -21°C. PCR was performed by adding Taq PCR master mix (Qiagen, 201443) and primer pair with overhanging adaptors compatible with the Iontorrent PGM platform (LGC Biosearch Technologies). All primers and single-stranded DNA-oligos are collected in ref. (Kristensen et al., 2024). PCR was performed under the following conditions: 95 °C 10 min; 36 cycles: 95 °C 30 s, 60 °C 45 s, 72 °C 30 s; and 72 °C 4 min, and the product was subsequently visualized using 2% Agarose gels (Invitrogen, G601802) confirming bands of expected sized (∼200 bps). Amplified products were then pooled and purified using QIAquick PCR purification kit (Qiagen, 28104) and a NanoDrop spectrophotometer to measure library cDNA concentration amplicon sequencing performed by PrimBio (USA) at a desired read coverage of 150-200 reads per barcode.

### Processing of sequencing data

FASTQ files were uploaded to Barracoda 1.8 (https://services.healthtech.dtu.dk/service.php?Barracoda-1.8). See details in (Bentzen et al., 2016). In brief, Barracoda first maps all reads to the constant primer and annealing regions followed by exclusion of incompletely mapped reads. Akey/SampleIDs are then assigned, and UMIs concatenated and counted as one followed by clonal reduction of UMI identities for each sampleID. Log fold change p-values indicating enrichment and statistical significance respectively were then calculated based on EdgeR comparing sorted products to triplicate baseline values from amplified multimer pools originating from aliquots taking and frozen before T cell staining with the multimer pool. P-values were corrected using the Benjamini-Hochberg procedure. Specific barcodes with FDR < 0.1% were defined as significant. All significant hits was represented with at least 1/1000 total reads to be defined as significant; a threshold used to exclude hits due to low coverage in input samples.

### Specific expansion using peptide pools

Specific expansion using peptide pools was implemented from ref. (Eberhardt et al., 2021) with the following changes. In short, two peptide pools were made covering laEBV / CEF antigens (77 peptides) and neuroantigens (75 peptides). 2.5 million PBMCs were then put into culture with either DMSO control, virus-derived or neuroantigen-derived peptides. Each peptide in the peptide pool was present at 1 µM. Short term cultures were kept in X-vivo 15 (LZ-BE02-060Q, Lonza), 5% human serum (H3667, Sigma-Aldrich), 50 IU/mL rIL2 (200-02, Peprotech), 25 ng/mL rIL7 (200-07, Peprotech), and 25 ng/mL rIL15 (200-15, Peprotech) for 13 days followed by washing and resting in cytokine free media overnight. Cultures were fed and/or split on a per need basis but typically on day 4 and every second day thereafter. Day 14 cultures were stained with CD3-BUV395, CD4-BUV563, CD8-BV480, antibody clones and cat numbers are found under antibody mastermix #2, as well as 1 PE-labelled and 1 APC-labelled dextran multimers. To stain for 20 specificities, expanded cultures were then split into 10 portions with the aim of staining 500.000-750.000 cells. PE- and APC-labeled pMHC dextran multimers were assembled as described under the previous section titled “Assembly of DNA-barcode labelled pMHC multimers”. The final staining concentration per multimer was increased to 5 µl / 60 µl stain volume to saturate T-cell binding to pMHC dextran multimers.

### Statistical analysis

All data analysis was performed in R version 4.0.5 with cowplot, ggpubr, ggbeeswarm, rstatix, and tidyverse (Wickham et al., 2019). Statistical analysis was performed in a university-hosted webservice Barracoda 1.8 (https://services.healthtech.dtu.dk/service.php?Barracoda-1.8) utilizing EdgeR (Robinson et al., 2009). All p-values for comparisons between exactly two samples were performed using non-parametric two-sample Wilcoxon test with alpha values set to 0.05, 0.01, 0.001, and 0.0001 for *, **, *** and ****, respectively. P-values for comparisons among multiple groups were calculated either by using Kruskal-Wallis test or by Dunn’s test except when a paired analysis was necessary in which case a paired pairwise Wilcox test was used. We adjusted p-values to control FDR at 0.05, when multiple p-values were generated with the same method within the same group and specified the resulting adjusted p-values as q-values. A two-sided Fischer’s exact test was performed to estimate significant enrichment of epitope recognition prevalence in one group over another. Paired Wilcoxon test was used for paired analyses.

## AUTHOR CONTRIBUTIONS

Conceptualization: N.P.K, N.W.E, F.S, S.R.H. Data curation: N.P.K, N.W.E, L.F.V, M.R.v.E. Formal analysis: N.P.K, S.R.H. Funding acquisition: F.S, S.R.H. Investigation: N.P.K, N.W.E, L.F.V, M.R.v.E. Methodology: A.K.B, T.T. Project administration: S.R.H. Resources: T.T, A.K.B, S.R.H. Supervision: S.R.H. Validation: N.P.K, N.W.E, L.F.V. Visualization: N.P.K. Writing original draft: N.P.K, S.R.H, L.F.V. Review and editing: all authors.

## ACKNOWLEDGMENTS

We thank all donors and patients for participating in the study. We thank the excellent technical assistance from B. Rotbøl, A.F. Løye, and A.D. Burkal for excellent technical assistance in handling blood samples and maintaining our instruments. The project was supported by, The Lundbeck Foundation, Fellowship R190-2014-4178 and the European Research Council ERC Consolidator Grant MIMIC 101045517 to S.R.H. The project was additionally supported by Scleroseforeningen, Denmark grant no. A44005 to SRH and LFV.

F.S holds a professorship at the Department of Clinical Medicine, University of Copenhagen sponsored by the Danish Multiple Sclerosis Society.

## CONFLICTS OF INTEREST

N.P.K and M.R.v.E have received speaker honoraria from Merck. SRH and AKB are coinventors of patents WO2015185067 (Determining antigen recognition through barcoding of MHC multimers) and WO2015188839 (General detection and isolation of specific cells by binding of labelled molecules) for the barcoded MHC technology that is licensed to Immudex. F.S has served on scientific advisory boards for, served as consultant for, received support for congress participation or received speaker honoraria from Biogen, Lundbeck, Merck, Neuraxpharm, Novartis, Roche and Sanofi. His laboratory has received research support from Biogen, Merck, Novartis, Roche and Sanofi. Finn Sellebjerg is section editor on Multiple Sclerosis and Related Disorders.

