## Supplementary material for "Mapping of CD8 T-cell recognition to latent EBV infection and neuroantigens reveals HLA-specific depletion of T-cell responses in multiple sclerosis"

**Supplemental table 1: HLA distribution**

| | HD | RRMS | PMS | p-value‡ |
|---|---|---|---|---|
| A*01:01 (yes / no) | 13 / 7 | 9 / 10 | 14 / 5 | ns |
| A*02:01 (yes / no) | 13 / 7 | 9 / 10 | 12 / 7 | ns |
| B*07:02 (yes / no) | 5 / 15 | 10 / 9 | 11 / 8 | ns ($p_{HDvPMS}$ = 0.054) |
| B*08:01 (yes / no) | 11 / 9 | 8 / 11 | 9 / 10 | ns |
| DRB1*15:01 (yes / no) | 1 / 19 | 13 / 6 | 11 / 8 | $p_{HDvPMS}$ = 3.9e-5<br>$p_{HDvRRMS}$ = 4.3e-4<br>$p_{PMSvRRMS}$ = 0.74 |

‡*A pairwise fischer's exact test was used for assessing HLA count differences between cohort groups.*

**Supplemental table 2: Control peptides**

| ID | Sequence | HLA | Protein | Ag.grp | Ag.patho.grp |
|---|---|---|---|---|---|
| v9 | YSEHPTFTSQY | A0101 | pp65 | Control | CMV |
| v15 | VTEHDTLLY | A0101 | pp50 | Control | CMV |
| v19 | VSDGGPNLY | A0101 | BP | Control | FLU |
| v14 | TPRVTGGGAM | B0702 | pp65 | Control | CMV |
| v20 | RPHERNGFTV | B0702 | pp65 | Control | CMV |
| v17 | RAKFKQLL | B0801 | BZLF1 | Control | EBV |
| v22 | ELRRKMMYM | B0801 | IE1 | Control | CMV |
| v1 | ELRSRYWAI | B0801 | NP | Control | FLU |
| v6 | GLCTLVAML | A0201 | BMLF1 | Control | EBV |
| v16 | YVLDHLIVV | A0201 | BRLF1 | Control | EBV |
| v24 | VLEETSVML | A0201 | IE1 | Control | CMV |
| v13 | NLVPMVATV | A0201 | pp65 | Control | CMV |
| v3 | GILGFVFTL | A0201 | MP | Control | FLU |
| v41 | ILKEPVHGV | A0201 | POL | Control | HIV |

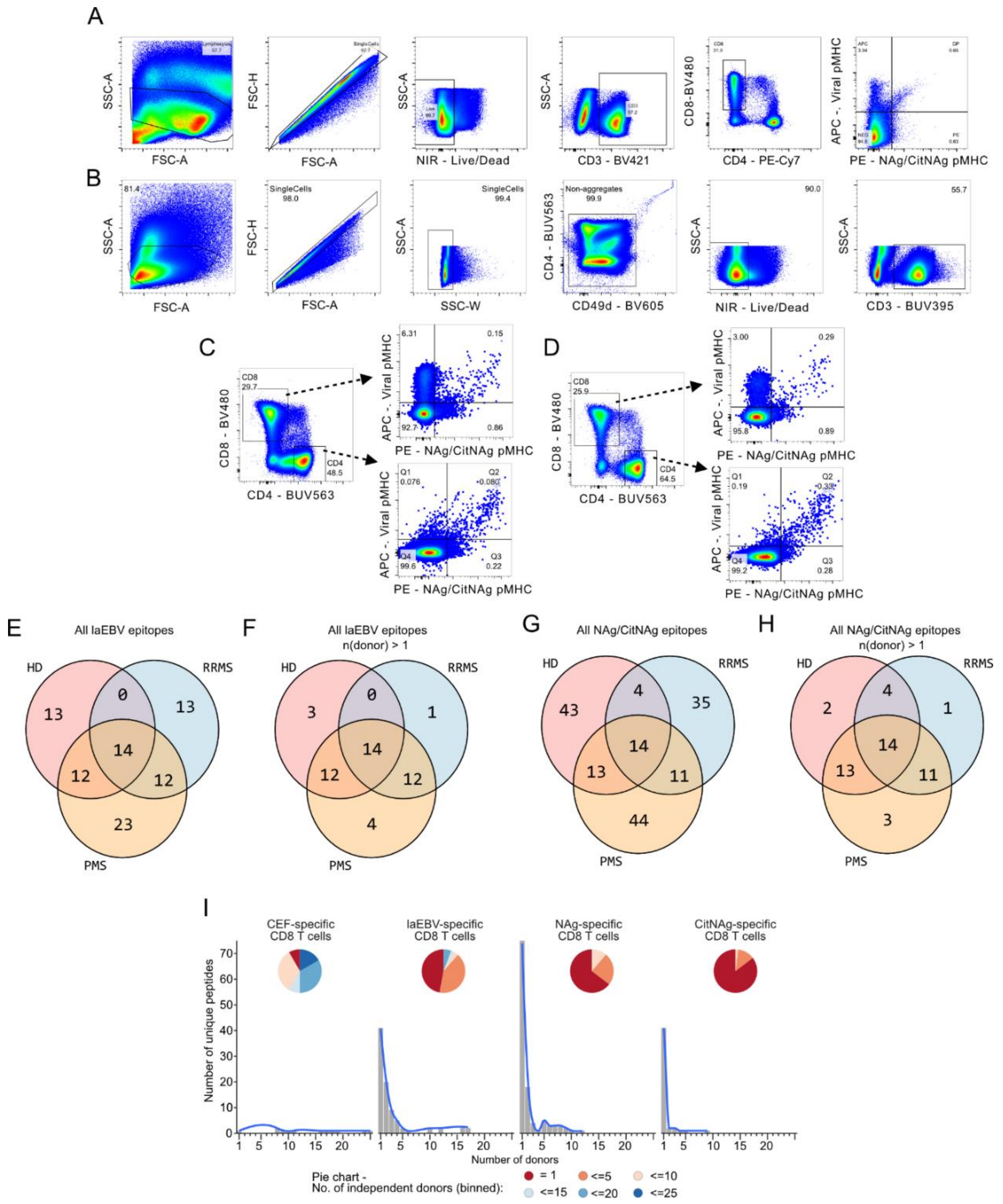


**Figure S1**: Flow cytometry gating. A) Gating strategy used for sorting multimer+ CD8 T cells from without additional phenotyping. B) Gating strategy used for sorting multimer+ CD8 T cells from donors with additional phenotypic antibodies included. Non-aggregates were not omitted during cell sorting and was only removed in post-sorting analysis steps. C-D) Two examples for multimer+ CD8 T cells binding CD4 and CD8 T cells. E) Venn diagram depicting laEBV epitopes discovered for each cohort group. F) Venn diagram as in F but only showing epitopes recognized in at least 2 donors. G) Venn diagram depicting NAg/CitNAg epitopes discovered for each cohort group. H) Venn diagram as in H but only showing epitopes recognized in at least 2 donors. I) Number of epitopes discovered with 1 or more independent observations in different donors across the cohort. Colored pie charts represents the relative distribution between epitopes of different numbers of binned independent donors.

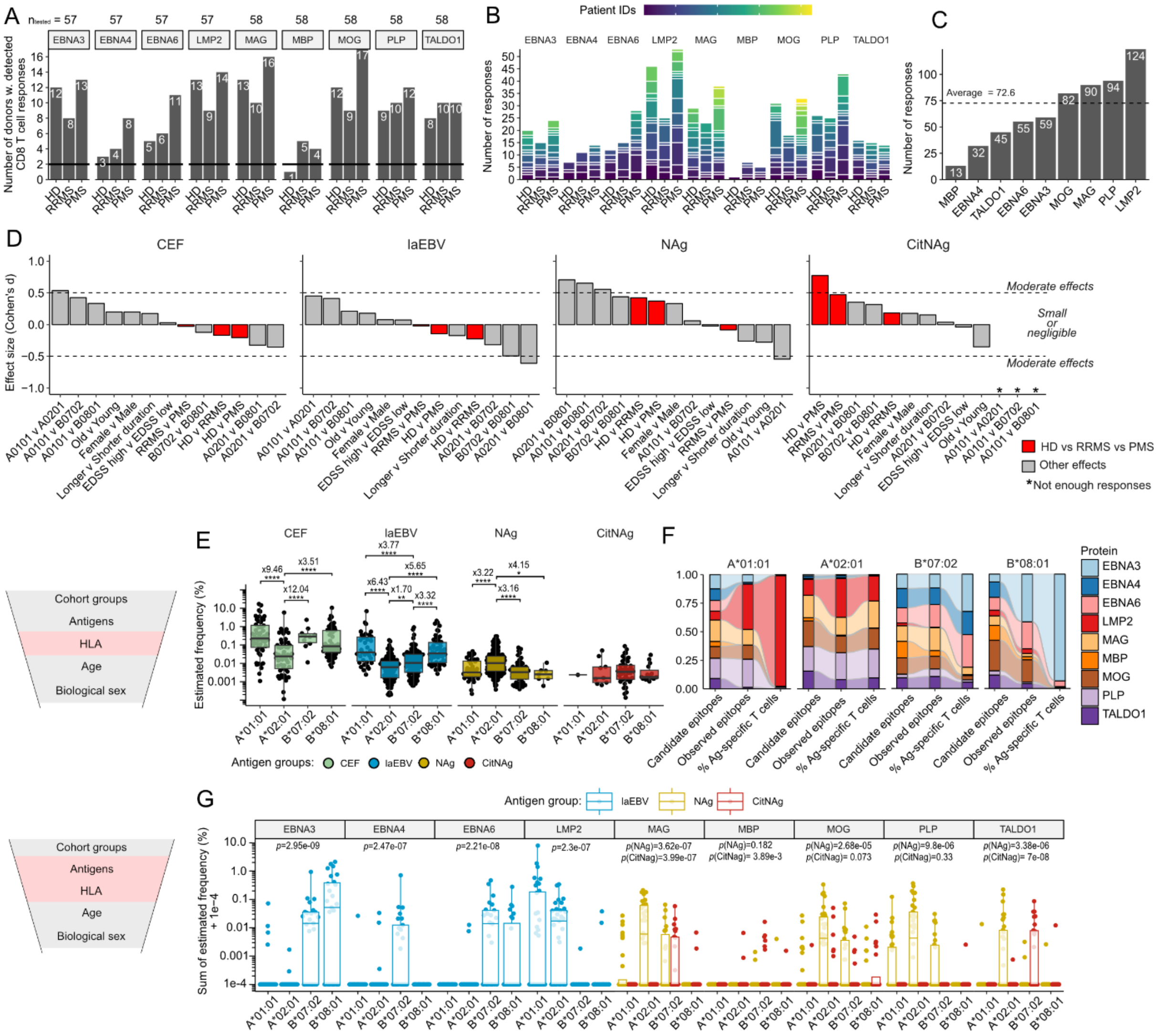

**Figure S2**: Supporting information on confounding factors in context of NAg- and IaEBV-specific immune recognition in MS patients and healthy donors. A) Number of independent donors that contribute data to each category of antigen-specific CD8 T cells across our cohort groups. Number of tested donors per antigen appears atop the plot. 1 donor was excluded from virus-specific analysis due to technical issues with an incompatible amplification primer. B) Similar to A, but with each colored segment representing a unique donor. C) Number of independent epitope-specific immune responses detected in in our cohort for each antigen of interest. D) Cohen's *d*, representing effect size of individual group-wise comparisons of antigen-specific cellular frequency. Dashed lines represent the conventional cut-off for small to moderate effect sizes (|d|≥0.5). Age, disease duration and EDSS scores were split by the overall median value with larger values being labeled as older, longer disease duration, and EDSS high. E) Comparison of read-based estimated multimer frequencies per HLA allele and antigen group. Fold change differences and p-values are indicated for each statistically significant comparison. F) Multimer and antigen distribution per HLA starting from an unweighted distribution of candidate epitopes, i.e. each multimer in our pMHC library is counted once, to the distribution of observed antigen-specific responses where each independent response is also counted once, to the observed antigen-specific response distribution weighted by the observed cellular frequency of the given antigen-specific response. G) Sum of estimated frequency on the y-axis is the total percentage of cells from a given donor that recognizes a given antigen. 0.0001 (1e-4) was added to avoid 0 values. Cohen's d values were corrected using Hedge's correction. P-values were calculated in E using Dunn's test and adjusted for multiple hypothesis testing using the Benjamini Hochberg approach.  P-values in G were calculated using a Kruskal-Wallis test. *, **, ***, and **** represented p-values lower than 0.05, 0.01, 0.001, and 0.0001, respectively. Near-significant values are indicated by numerical p-values rounded to the nearest second digit.

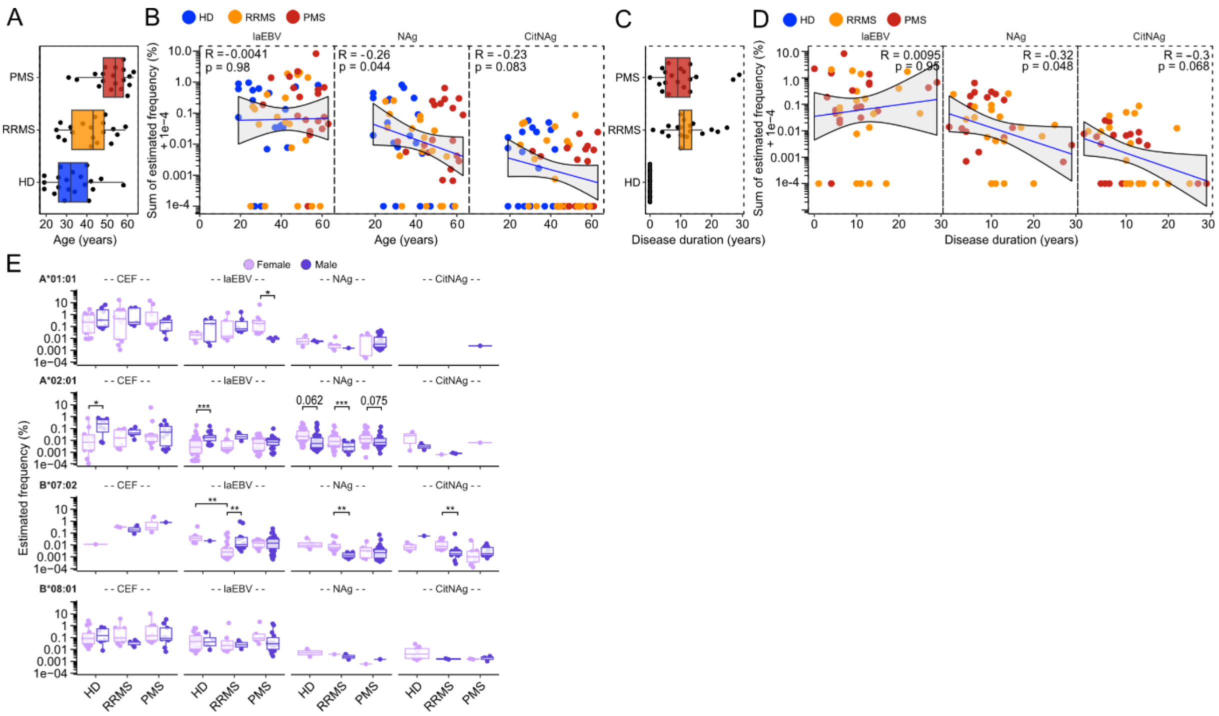


**Figure S3**: Supporting information on age and biological sex as possible confounders of antigen-specific frequency. A) Boxplot summary of age variability across our cohort. B) Spearman correlation and scatter plot of age and the total cellular frequency of antigen-specific immune responses per donor. C) Boxplot summary of disease duration variability across our cohort. D) Spearman correlation and scatter plot of disease duration and the total cellular frequency of antigen-specific immune responses per donor. E) Estimated frequency of laEBV and NAg/CitNAg-specific immune responses stratified by sex and HLA genotype. R and p-values were calculated by spearman correlation in B and D. P-values were calculated using a Dunn's test in E. *, **, and ***, represented p-values lower than 0.05, 0.01, and 0.001, respectively. Exact p-values were shown for near-significant comparisons.

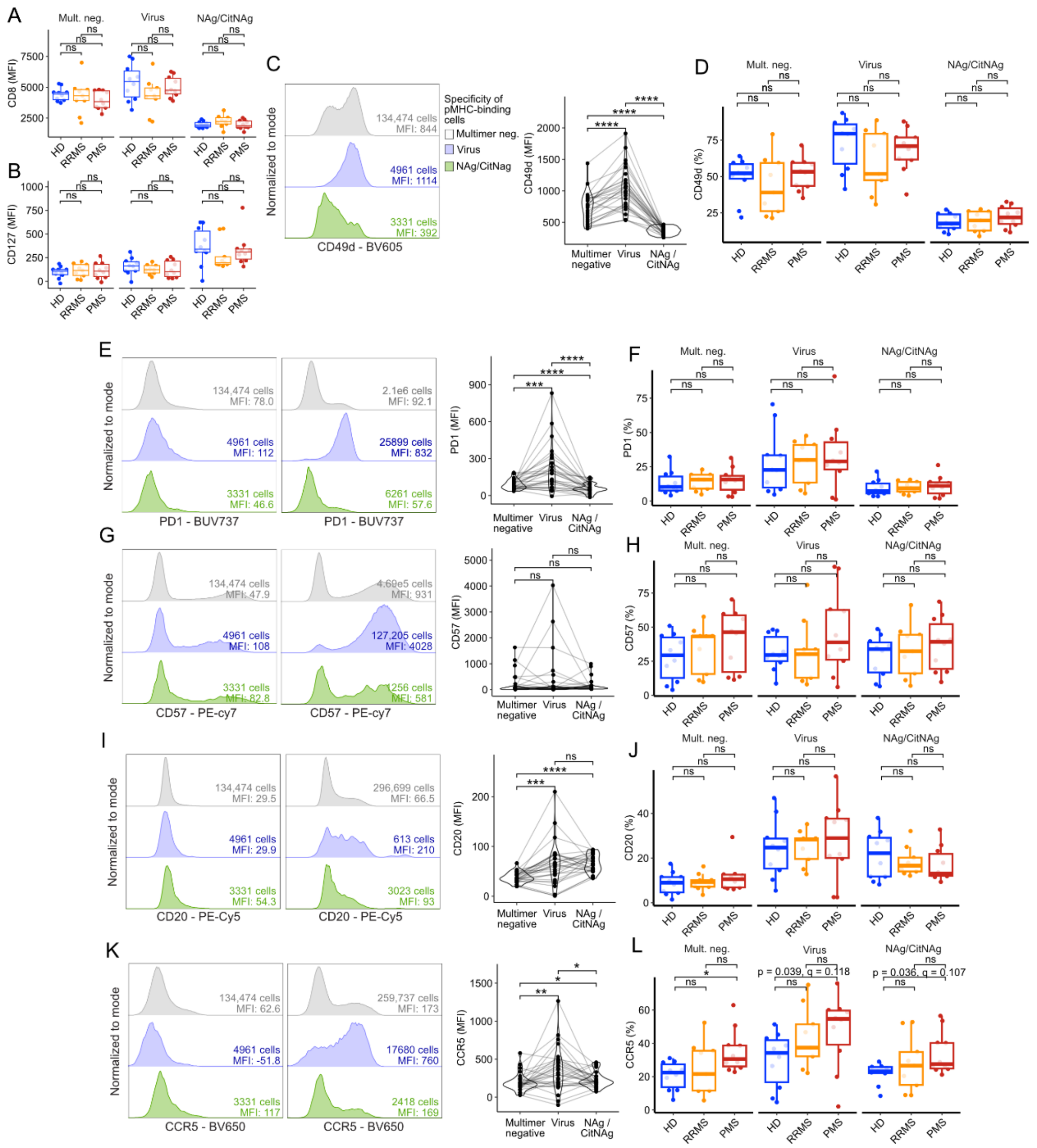


**Figure S4:** Supporting information on the immunophenotypes of discovered antigen-specific virus- and neuroantigen-recognizing CD8 T cells. A) CD8-BV480 MFI values for multimer negative, virus-specific and NAg/CitNAg-specific CD8 T cells. Values are grouped according to cohort groups. B) CD127-APC-R700 MFI values for multimer negative, virus-specific and NAg/CitNAg-specific CD8 T cells. Grouped as in (A). C) Left, representative CD49d-BV605 fluorescent intensities reported for multimer negative, multimer negative, virus-specific and NAg/CitNAg-specific CD8 T cells. Right, paired analysis of MFI values compared between cohort groups. D) CD49d-BV605 MFI values for multimer negative, virus-specific and NAg/CitNAg-specific CD8 T cells. Grouped as in (A). E-F) Representative PD1-BUV737 fluorescent intensities and paired analysis followed by a comparison between cohort groups of PD1-BUV737 MFI values in (F). G-H) Representative CD57-PE-cy7 fluorescent intensities and paired analysis followed by a comparison between cohort groups of CD57-PE-cy7 MFI values in (H). I-J) Representative CD20-PE-cy5 fluorescent intensities and paired analysis followed by a comparison between cohort groups of CD20-PE-cy5MFI values in (J). K-L) Representative CCR5-BV650 fluorescent intensities and paired analysis followed by a comparison between cohort groups of CCR5-BV650 MFI values in (L). P-values were calculated using a paired Wilcox test for MFI values, and a Dunn‘s test for the remaining differences. All p-values were corrected using the Benjamini-Hochberg approach. *, **, ***, and **** represented p-values lower than 0.05, 0.01, 0.001, and 0.0001, respectively. Exact p-values were shown for near-statistically significant comparisons or in case of adjusted p-values (q) being non-significance, where the p-value was significant.

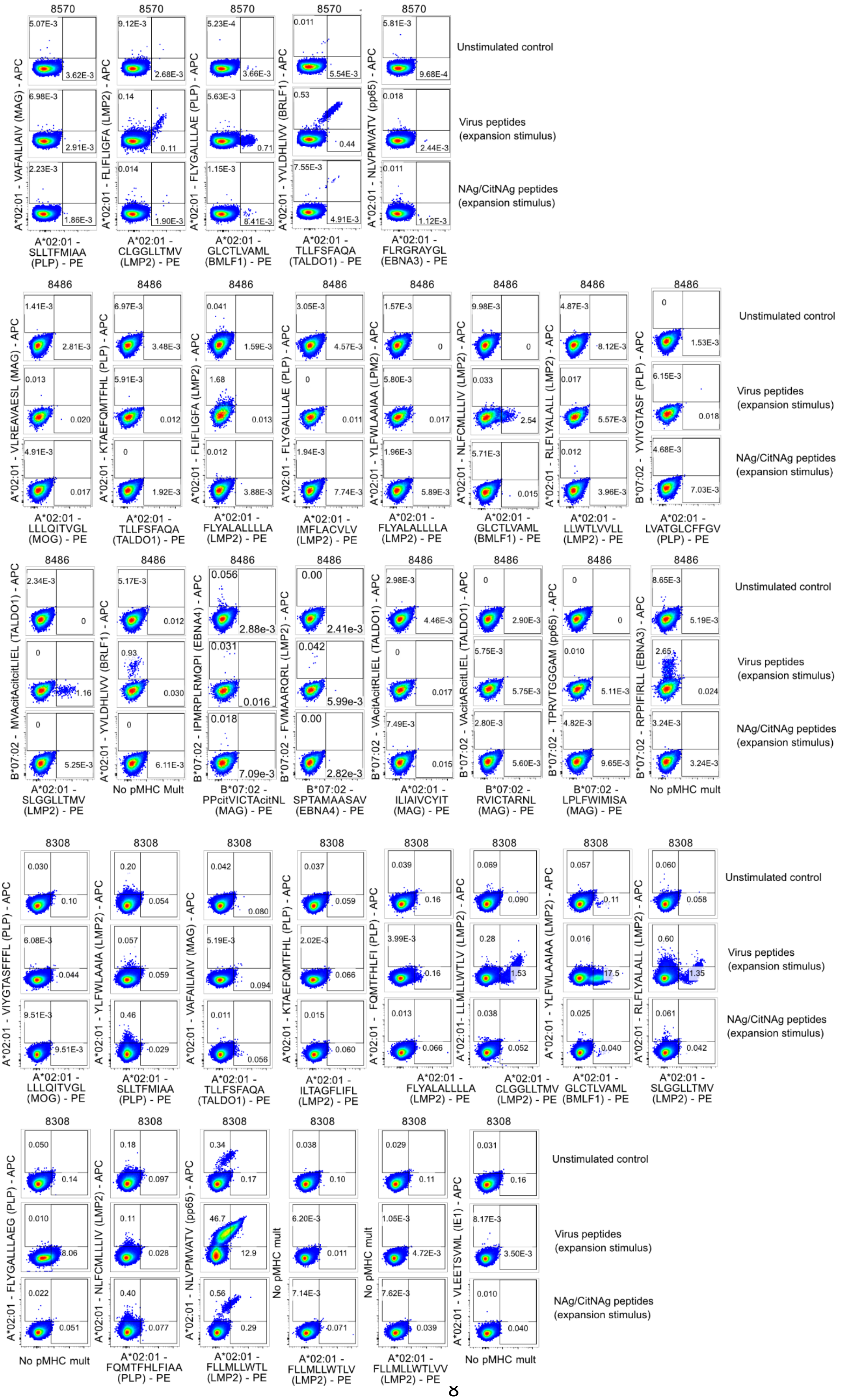
8570
Unstimulated control
Virus peptides (expansion stimulus)
NAg/CitNAg peptides (expansion stimulus)
A*02:01 - VAFAILIAIV (MAG) - APC
A*02:01 - FLFLIGFA (LMP2) - APC
A*02:01 - FLYGALLLAE (PLP) - APC
A*02:01 - YVLDHLIVV (BRLF1) - APC
A*02:01 - NLVPMVATV (pp65) - APC
A*02:01 - SLLTFMIAA (PLP) - PE
A*02:01 - CLGGLLTMV (LMP2) - PE
A*02:01 - GLCTLVAML (BMLF1) - PE
A*02:01 - TLLFSFAQA (TALDO1) - PE
A*02:01 - FLRGRAYGL (EBNA3) - PE
8486
A*02:01 - VLREAVAESL (MAG) - APC
A*02:01 - KTAEFQMTFHL (PLP) - APC
A*02:01 - FLFLIGFA (LMP2) - APC
A*02:01 - FLYGALLLAE (PLP) - APC
A*02:01 - YLFWLAAIAA (LPM2) - APC
A*02:01 - NLFCMLLLIV (LMP2) - APC
A*02:01 - RLFLYALALL (LMP2) - APC
B*07:02 - YVIYGTASF (PLP) - APC
A*02:01 - LLLQITVGL (MOG) - PE
A*02:01 - TLLFSFAQA (TALDO1) - PE
A*02:01 - FLYALALLLA (LMP2) - PE
A*02:01 - IMFLACVLV (LMP2) - PE
A*02:01 - FLYALALLLA (LMP2) - PE
A*02:01 - GLCTLVAML (BMLF1) - PE
A*02:01 - LLWTLVVLL (LMP2) - PE
A*02:01 - LVATGLCFFGV (PLP) - PE
B*07:02 - MVAcitAcitLIEL (TALDO1) - APC
A*02:01 - YVLDHLIVV (BRLF1) - APC
B*07:02 - IPMRPLRMQPI (EBNA4) - APC
B*07:02 - FVMAARQRL (LMP2) - APC
B*07:02 - VAcitRLIEL (TALDO1) - APC
B*07:02 - VAcitARcitLIEL (TALDO1) - APC
B*07:02 - TPRVTGGGAM (pp65) - APC
B*07:02 - RPPIFIRLL (EBNA3) - APC
A*02:01 - SLGGLLTMV (LMP2) - PE
No pMHC Mult
B*07:02 - PPcitVICTAcitNL (MAG) - PE
B*07:02 - SPTAMAASAV (EBNA4) - PE
A*02:01 - ILIAIVCYIT (MAG) - PE
B*07:02 - RVICTARNL (MAG) - PE
B*07:02 - LPLFWIMISA (MAG) - PE
No pMHC mult
8308
A*02:01 - VIYGTASFFFL (PLP) - APC
A*02:01 - YLFWLAAIA (LMP2) - APC
A*02:01 - VAFAILIAIV (MAG) - APC
A*02:01 - KTAEFQMTFHL (PLP) - APC
A*02:01 - FQMTFHLFI (PLP) - APC
A*02:01- LLMLWTLV (LMP2) - APC
A*02:01 - YLFWLAAIAA (LMP2) - APC
A*02:01 - RLFLYALALL (LMP2) - APC
A*02:01 - LLLQITVGL (MOG) - PE
A*02:01 - SLLTFMIAA (PLP) - PE
A*02:01 - TLLFSFAQA (TALDO1) - PE
A*02:01 - ILTAGFLIFL (LMP2) - PE
A*02:01 - FLYALALLLA (LMP2) - PE
A*02:01 - CLGGLLTMV (LMP2) - PE
A*02:01 - GLCTLVAML (BMLF1) - PE
A*02:01 - SLGGLLTMV (LMP2) - PE
A*02:01 - FLYGALLAEG (PLP) - APC
A*02:01 - NLFCMLLLIV (LMP2) - APC
A*02:01 - NLVPMVATV (pp65) - APC
No pMHC mult
A*02:01 - VLEETSVML (IE1) - APC
No pMHC mult
A*02:01 - FQMTFHLFIAA (PLP) - PE
A*02:01 - FLLMLWTL (LMP2) - PE
A*02:01 - FLLMLWTLV (LMP2) - PE
A*02:01 - FLLMLWTLVV (LMP2) - PE
No pMHC mult

**Figure S5:** Raw flow plots for multimer+ CD8+ T cells following 14 days of expansion from peptide-stimulated PBMCs. Cells were pregated on live, CD3+, CD4-, CD8+ T cells. Cases with cross-binding populations were considered technical artefacts likely due to high differences in peptide-binding affinity to MHC.

**Supplemental table 3: Prevalence of recognition data for all peptides tested**

- See separately attached file
